# FBXW7 regulates MYRF levels to control myelin capacity and homeostasis in the adult CNS

**DOI:** 10.1101/2024.10.15.618515

**Authors:** Hannah Y. Collins, Ryan A. Doan, Jiaxing Li, Jason E. Early, Megan E. Madden, Tyrell Simkins, David A. Lyons, Kelly R. Monk, Ben Emery

## Abstract

Myelin, along with the oligodendrocytes (OLs) that produce it, is essential for proper central nervous system (CNS) function in vertebrates. Although the accurate targeting of myelin to axons and its maintenance are critical for CNS performance, the molecular pathways that regulate these processes remain poorly understood. Through a combination of zebrafish genetics, mouse models, and primary OL cultures, we found FBXW7, a recognition subunit of an E3 ubiquitin ligase complex, is a regulator of adult myelination in the CNS. Loss of *Fbxw7* in myelinating OLs resulted in increased myelin sheath lengths with no change in myelin thickness. As the animals aged, they developed progressive abnormalities including myelin outfolds, disrupted paranodal organization, and ectopic ensheathment of neuronal cell bodies with myelin. Through biochemical studies we found that FBXW7 directly binds and degrades the N-terminal of Myelin Regulatory Factor (N-MYRF), to control the balance between oligodendrocyte myelin growth and homeostasis.

## Introduction

In the vertebrate central nervous system (CNS), myelin is produced by the specialized glial cells called oligodendrocytes (OLs). OLs wrap segments of axons, creating a multi-layered sheath that speeds the transmission of nerve impulses and provides critical support to axons^1–4^. The formation and targeting of myelin is influenced by external cues such as axonal caliber and neuronal activity, but is also tightly controlled by cell intrinsic programs^5–8^. As oligodendrocyte precursor cells (OPCs) differentiate from a dynamic and proliferative state into a relatively stable post-mitotic myelinating OL, they reorganize and expand their cytoskeleton, cytoplasm, and membrane^9–11^, requiring significant transcriptional changes^12–16^. Although OLs are long-lived, surviving up to years in mice and decades in human white matter tracts, the myelin constituents themselves turn over comparatively rapidly, with a half-life of months^17–20^. Individual myelin sheaths can also be remodeled throughout life^8,21,22^. Therefore, understanding the molecular pathways and mechanisms that balance myelin growth and homeostasis is crucial for understanding myelin’s role in health, aging, and disease.

Previous work from our lab and others has identified F-box and WD repeat domain-containing protein 7 (FBXW7) as a key negative regulator of developmental myelination by both Schwann cells in the peripheral nervous system (PNS) and OLs in the CNS of zebrafish^23–26^. Within the PNS, FBXW7 regulates Schwann cell numbers, along with their myelin thickness^25^. Surprisingly, loss of *Fbxw7* also results in a breakdown of the normal 1:1 relationship between myelinating Schwann cells and axons, with individual *Fbxw7* conditional knockout Schwann cells aberrantly myelinating multiple axons^25^. Within the CNS of zebrafish, *fbxw7* regulates neural stem cell fate through Notch signaling, biasing the cells towards an OPC fate and increasing the pool of OL lineage cells in the spinal cord^23^. At later stages, loss of *fbxw7* leads to increased myelin sheath length, attributed to dysregulation of mTOR signaling^24^. *Fbxw7* encodes the F-box domain containing recognition subunit of a SKP1-Cullin-Fbox (SCF) E3 ubiquitin ligase complex. It mediates its biological effects through targeting specific proteins for proteasomal degradation, thus controlling their total levels in the cell^23,24,27–29^. FBXW7 substrates are highly variable between cell types and have not yet been investigated in myelinating cells in an unbiased manner.

To interrogate the role of FBXW7 in the regulation of CNS myelination, we used a combination of zebrafish, primary mammalian OL cultures, and conditional knockout mouse models. We found that inactivation of *fbxw7* in developing zebrafish resulted in enhanced OL maturation in the spinal cord. Strikingly, conditional ablation of the *Fbxw7* gene in mature OLs in the adult mouse CNS also increased myelin sheaths but also resulted in progressive myelin abnormalities including outfolds, disrupted paranodal organization, and ectopic ensheathment of neuronal cell bodies with myelin. We found that *Fbxw7* deficient OLs had no changes in mTOR protein levels in primary mammalian OL cultures, suggesting the myelin phenotypes were not a consequence of dysregulated mTOR signaling. Previous work in hepatocarcinoma cells identified the pro-myelination transcription factor Myelin Regulatory Factor (MYRF) as a target of FBXW7^30^. We demonstrate that the N-terminus of MYRF is a direct target of FBXW7 in OLs both *in vitro* and *in vivo*, with levels of N-MYRF protein and many of its transcriptional targets substantially increased in *Fbxw7* deficient OLs. We also find that *myrf* haploinsufficiency is sufficient to reverse both the increase in OL numbers and myelination seen in *fbxw7* knockout fish. Taken together, our findings demonstrate that FBXW7 is an evolutionarily conserved negative regulator of OL myelination and that its negative regulation of MYRF in the adult CNS is required for long-term myelin homeostatic maintenance.

## Results

### Fbxw7 regulates OPC specification and OL myelination

In zebrafish, global *fbxw7* mutations cause hypermyelination in both the PNS and CNS^23–25^. Previously published zebrafish *fbxw7* mutant alleles are late mutations within the region of the gene encoding the WD40 substrate recognition domain of the FBXW7 protein; therefore, existing mutants may not be complete loss-of-function alleles^23,26^. We used CRISPR-Cas9-mediated genome editing to create a new mutation, *fbxw7^vo86^*. This mutation introduces a frameshift and early stop codon in exon 5, which encodes the F-Box domain that allows FBXW7 to interact with its E3 complex (Supplementary Fig. 1a, b). Through *in situ* hybridization, live imaging, and qPCR, we found that the *fbxw7^vo86^* mutation phenocopies the previously described N-ethyl-N-nitrosourea (ENU)-generated *fbxw^stl64^* mutants, including increased *myelin basic protein* (*mbp*) RNA levels and myelin intensity in the dorsal spinal cord (Fig. 1a, b and Supplemental Fig. 1c-e). Since *fbxw7^vo86^* phenocopies the original ENU-generated mutation, we concluded these mutations both represent full loss-of-function alleles. As the *fbxw7^vo86^*mutation disrupts the F-Box domain, the same region targeted in the Fbxw7^fl/fl^ mouse line^31^ also used in our studies (described below), this line was used for all subsequent analyses.

**Fig. 1.**
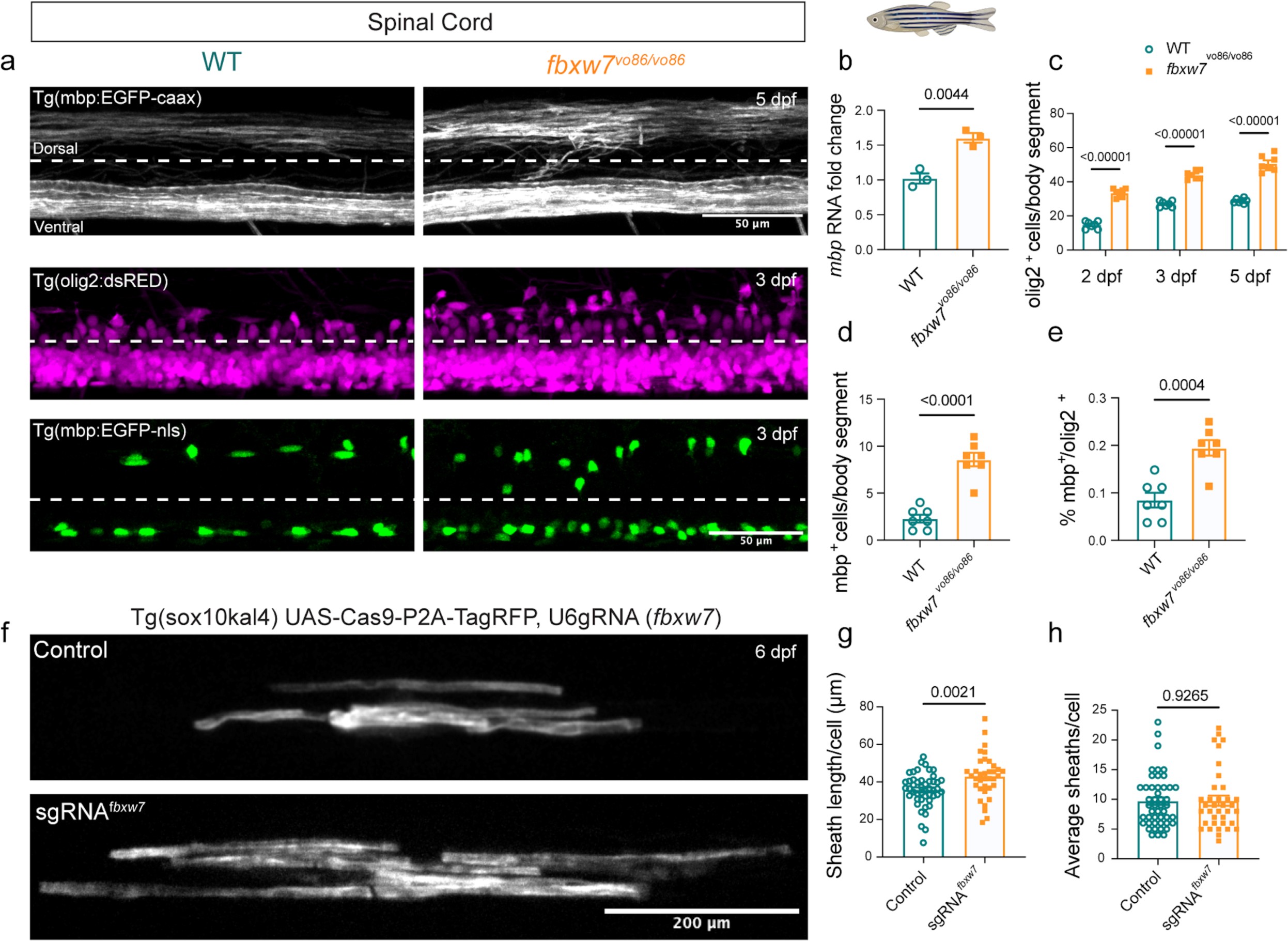
Fbxw7 regulates OPC specification and OL myelination in the zebrafish spinal cord. **a** Spinal cords of *fbxw7*^vo86^ and WT controls in (top) *Tg(mbp:eGFP-caax)*, (middle) *Tg*(*olig2:dsRED^+^),* and (bottom) *Tg(mbp:nls-eGFP)* transgenic backgrounds at 3 and 5 days post-fertilization (dpf; dotted-line delineates dorsal and ventral spinal cord tracts). **b** *mbp* RNA levels evaluated by qRT-PCR in 5 dpf *fbxw7*^vo86^ and WT control whole larvae. Average ± SEM, N = 3 (larvae). Statistical significance determined by unpaired, two-tailed Student’s t test. **c** Quantification of *olig2:dsRED^+^* cells/body segment in the dorsal spinal cord at 2, 3, and 5 dpf. Average ± SEM, N = 7 (larvae). Statistical significance determined by two-way ANOVA. **d** Quantification of *mbp:nls-eGFP^+^*OLs in the dorsal spinal cord/body segment at 3 dpf. Average ± SEM, N = 7 (larvae). Statistical significance determined by unpaired, two-tailed Student’s t test. **e** *mbp-nls:eGFP^+^* OLs normalized to total number of *olig2:dsRED^+^*cells/body segments in dorsal spinal cord at 3 dpf. Average ± SEM, statistical significance determined by unpaired, two-tailed Student’s t test. **f** Representative images of individual labelled OLs from cell-type specific CRISPR-Cas9 knock-down of *fbxw7* in *sox10* expressing cells in the zebrafish spinal cord at 6 dpf. The UAS-nlscas9-P2A-TagRFPT-caax construct allows visualization of cas9 expressing cells. **g** Quantification of average sheath length and **h** number per OL in controls and with *fbxw7* knockdown. Average ± SEM, Control N = 47 (cells), sgRNA*^fbxw7^* N=36 (cells). Statistical significance determined by unpaired, two-tailed Student’s t test. Created in BioRender. Emery, B. (2024) BioRender.com/o45h010.

In prior work, global disruption of *fbxw7* in zebrafish led to enhanced OPC specification through disinhibition of Notch signaling, observed as an increase in *mbp* expression and numbers of *olig2:dsRED*-expressing cells^23^. Consistent with this, *fbxw7^vo86^* mutants present with a significant increase in *mbp:EGFP-caax* expression and *olig2:dsRED^+^* cell numbers in the developing dorsal spinal cord relative to wild-type controls at 2-, 3-, and 5 days post-fertilization (dpf) (Fig. 1a-c). To examine later stages of the OL lineage, we crossed *fbxw7^vo86^* mutants into a transgenic *Tg(mbp-nls:eGFP)* line to label mature OL nuclei. Relative to wild-type clutchmates, the density of *mbp-nls:eGFP-*expressing cells was significantly increased in *fbxw7^vo86^* mutants, even when normalized to the increased numbers of OL lineage cells (*olig2:dsRED^+^*) at 3 dpf (Fig. 1a, d, e). To determine whether *fbxw7* could regulate myelination via an OL intrinsic mechanism, we utilized a cell-specific CRISPR-Cas9-mediated gene disruption system our lab had previously developed^32^. We found that *sox10*-driven disruption of *fbxw7* in OL lineage cells resulted in a significant increase in myelin sheath lengths at 6 dpf, with no change in the number of sheaths formed per individual OL (Fig. 1f-h). These data suggest that *fbxw7* regulates OL myelination though a cell-autonomous mechanism in the developing zebrafish spinal cord.

### FBXW7 regulates OL myelin sheath length, paranodal organization, and myelin homeostasis in both grey and white matter

Our findings in zebrafish indicated that *fbxw7* regulates key aspects of myelin growth early in development. Whether this role is conserved in mammalian systems and whether Fbxw7 regulates myelination past these early developmental stages has not been explored to date. To address this, we created an inducible *Fbxw7* knockout (icKO) mouse by crossing Fbxw7^fl/fl^ mice^31^ to the Plp1-CreERT line^33^, allowing for tamoxifen (TAM)-inducible knockout of *Fbxw7* in mature OLs. Fbxw7^fl/fl^; Plp1-CreERT^+^ mice (Fbxw7^ΔPlp1^) and their CreERT negative littermate controls (Fbxw7^fl/fl^) were treated with TAM at 8 weeks of age, and tissue was taken 1-, 3-, and 6-months post-TAM for subsequent analyses (Fig. 2a).

**Fig. 2.**
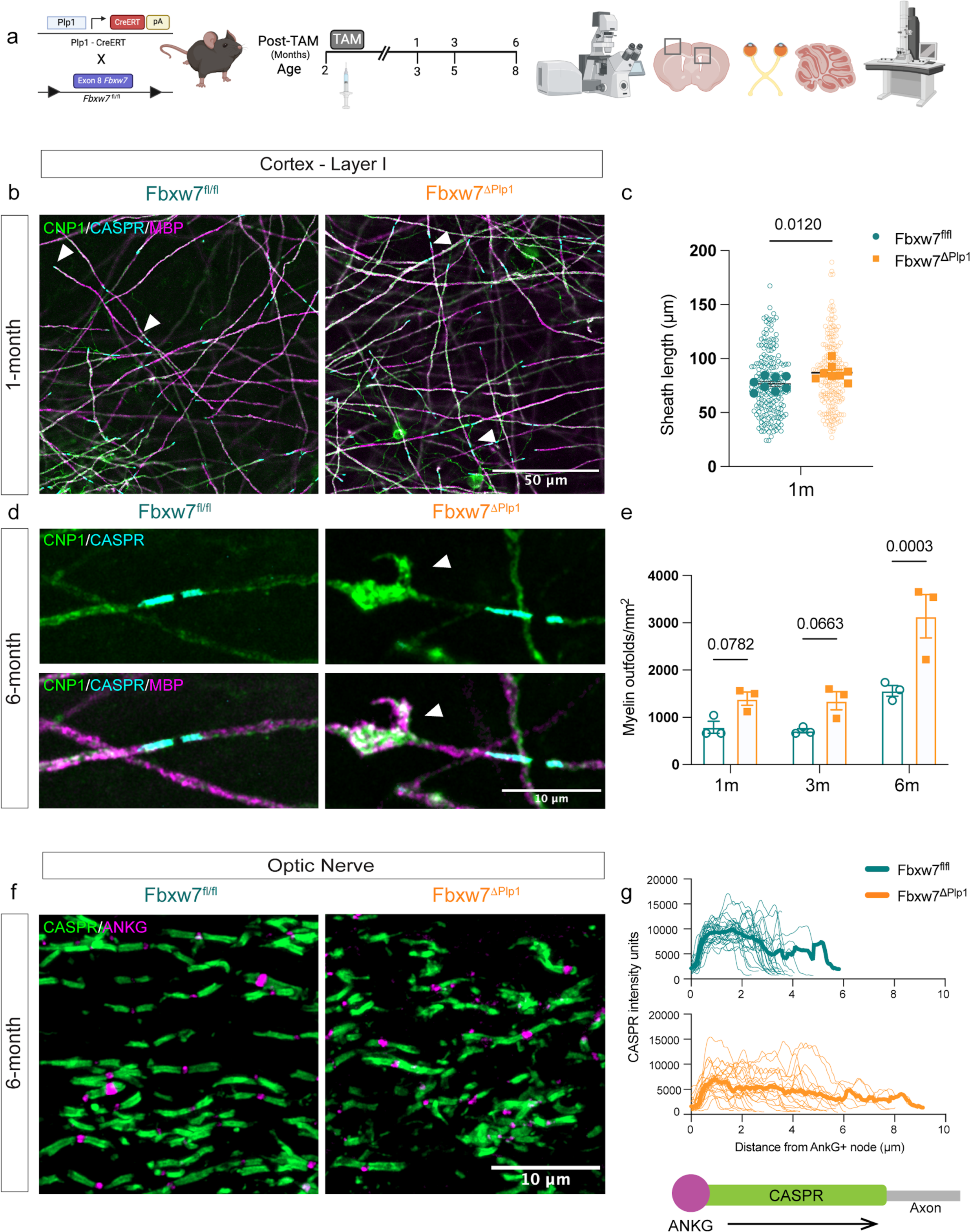
F*b*xw7 regulates OL internode length, myelin homeostasis, and paranode organization in mouse grey and white matter. **a** Schematic of Fbxw7^fl/fl^; Plp1-CreERT (Fbxw7^ΔiPlp1^) mouse line and experimental pipeline. **b** Representative images of myelin internodes in Fbxw7^fl/fl^ and Fbxw7^ΔPlp1^ 1 months post-tamoxifen (TAM) in mouse layer I of the primary somatosensory cortex (pSS), stained with MBP, CNP1, and CASPR. Arrowheads show CASPR^+^ boundaries of internodes. **c** Quantification of sheath length from layer I pSS. Colored bolded dots represent average of ROI from individual biological replicates Fbxw7^fl/fl^ N=4, Fbxw7^ΔPlp1^ N=4 (mice), small hollow dots represent values from individual sheaths (Fbxw7^fl/fl^ N=197, Fbxw7^ΔPlp1^ N=210). Statistical significance determined by unpaired, two-tailed Student’s t-test on ROI averages. **d** High magnification images of myelin outfolds (arrowheads) in pSS cortex 6 months post-TAM. **e** Quantification of myelin outfolds in the pSS. Biological replicates as per b and c. **f** Representative images of CASPR and ANKG stained longitudinal optic nerve sections from Fbxw7^fl/fl^ and Fbxw7^ΔPlp1^ mice at 6 months post-TAM. **g** Quantification of CASPR straining intensity as a function of distance from ANKG^+^ node. Thick lines represent median for the genotype, thin lines represent individual heminode intensity histograms. Fbxw7^fl/fl^ N=4 (total nodes= 27), Fbxw7^ΔPlp1^ N=4 (total nodes=32) from 3 technical replicates (images). Created in BioRender. Emery, B. (2024) BioRender.com/d36g650.

To determine if loss of *Fbxw7* in mature OLs regulated myelin sheath maintenance, we sectioned cortical flat-mounts of layer I and performed immunofluorescence (IF) for MBP (compact myelin), 2’,3’-cyclic nucleotide 3’ phosphodiesterase (CNP1, non-compact myelin), and contactin-associated protein (CASPR; paranodes). At 1-month post-TAM, Fbxw7^ΔPlp1^ OLs showed a significant increase in myelin sheath length in the primary somatosensory cortex (pSS) (Fig. 2b, c), indicating that FBXW7 regulates myelin capacity in mature OLs. Additionally, we observed an increase in the number of MBP/CNP1^+^ focal hyperintensities that appeared to be myelin outfolds, which increased significantly as the animals aged (Fig. 2d, e). To investigate FBXW7’s role in white matter, we performed IF on Fbxw7^ΔPlp1^ optic nerves at 6-months post-TAM. We found a pronounced breakdown of nodal organization with a significant broadening in the distribution of CASPR intensity and length at each heminode. (Fig. 2f, g). While Fbxw7^ΔPlp1^ animals had no change in weight or general health (Supplemental Fig. 2a), we observed a significant reduction of OL numbers in the upper cortex (layers 1-4) at 6 months post-TAM in Fbxw7^ΔPlp1^ animals relative to their age-matched control littermates (Supplemental Fig. 2b). Interestingly, this reduction was only observed above layer IV, with no change in OL numbers in deeper cortical layers or change in astrocyte reactivity (Supplemental Fig. 2c, f). We found no change in OPC numbers at any of these timepoints or regions (Supplemental Fig. 2d, e). Collectively, these data suggest that FBXW7 functions in many aspects of OL biology, from early modulation of sheath lengths to long term maintenance of paranode organization and myelin homeostasis in mammalian OLs.

Given the optic nerves of Fbxw7^ΔPlp1^ animals showed evidence of outfolds and disrupted nodal organization at 6-months post-TAM by IF, we next wanted to assess the ultrastructure of the myelin. We therefore performed transmission electron microscopy (TEM) on optic nerves from Fbxw7^ΔPlp1^ and Fbxw7^fl/fl^ littermate controls 6 months post-TAM. Consistent with our observations in layer I of the cortex, we also found a significant increase in the number of myelin outfolds in Fbxw7^ΔPlp1^ optic nerves (Fig. 3a, b). While control animals did have outfolds at low frequencies, as expected at 8 months of age^22^, the number and average length of outfolds in the Fbxw7^ΔPlp1^ was significantly higher (Fig. 3b, c). Although myelin ultrastructure was disrupted in the optic nerve, we found no change in the proportion of axons myelinated or their corresponding g-ratios when severe outfolds were excluded from analyses (Fig. 3d, e). Along with outfolds, we also observed other myelin abnormalities throughout the optic nerve including myelin whorls and double myelin sheaths (sheaths enveloped by an overlying sheath) (Fig. 3f-h). This double myelination was also observed by IF in layer I of the pSS cortex, where we found CASPR^+^ paranodes under MBP^+^ myelin sheaths in Fbxw7^ΔPlp1^ mice, suggestive of double myelinated axons (Fig. 3i). Similarly, the outfolds in the white matter tracts of the opctic nerve and corpus callosum were so severe they were also visible at the light level in Fbxw7^ΔPlp1^ animals as hyperintense MBP+ puncta (Supplemental Fig. 3a-c). We found no change in the number of OLs or OPCs (Supplemental Fig. 3b-e). Taken together, these data show that FBXW7 is a conserved regulator of myelin sheath length, which was independent of myelin sheath thickness, as well as long-term maintenance of myelin homeostasis and nodal organization.

**Fig. 3.**
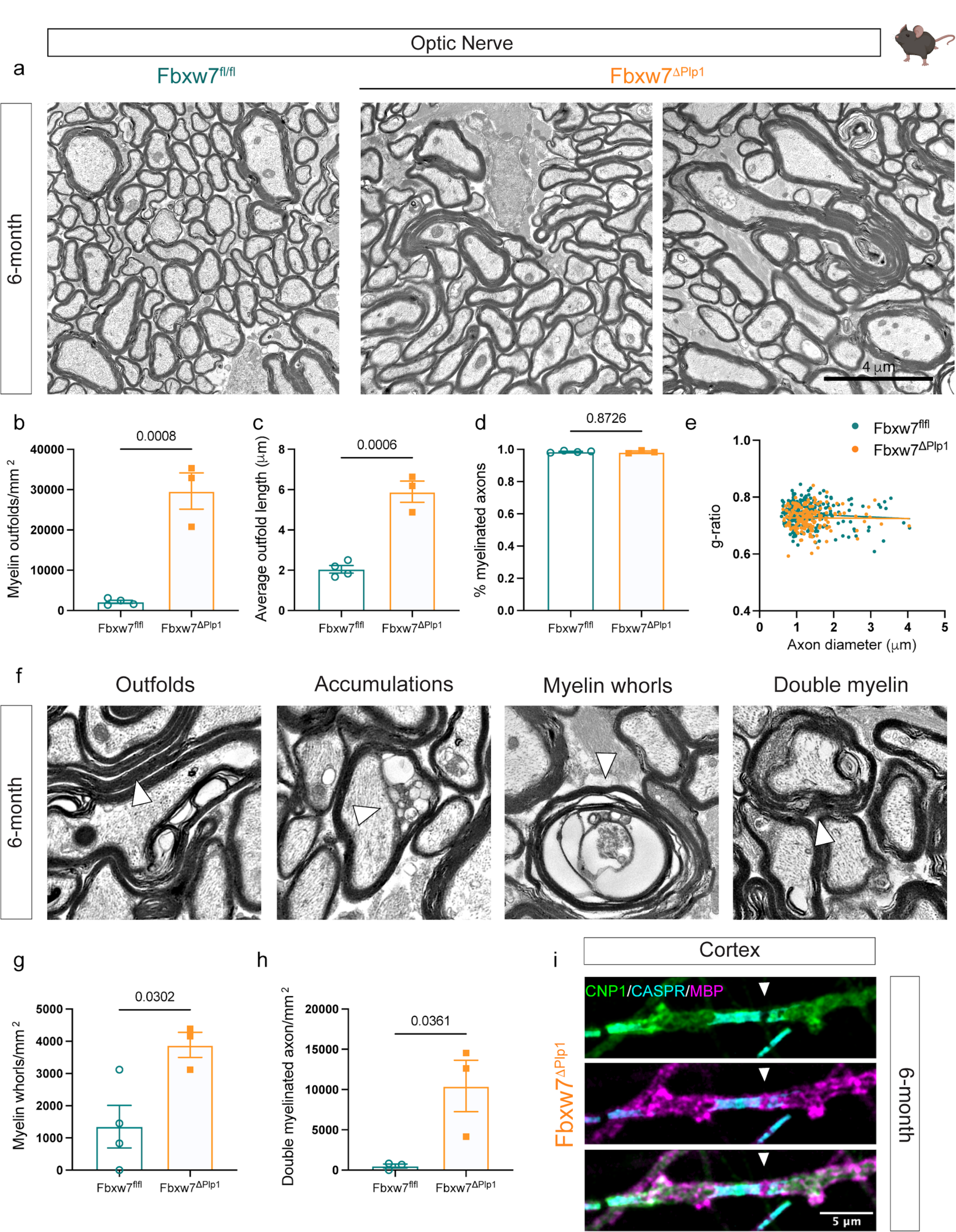
Loss of *Fbxw7* in OLs results in severe myelin outfolds in the optic nerve. **a** Representative Transmission Electron Microscopy (TEM) images of optic nerves from Fbxw7^fl/fl^ and Fbxw7^ΔPlp1^ mice at 6 months post-TAM. **b-e** Quantification of number of outfolds (**b)**, severity of outfolds (**c**), number of myelinated axons (**d**), and g-ratios (**e**). **f** Representative images of different myelin abnormalities in Fbxw7^ΔPlp1^ optic nerves at 6-months post-TAM. **g-h** Quantification of OL accumulations, myelin whorls, and double myelin. All data displayed as average ± SEM. Fbxw7^fl/fl^ N=4, Fbxw7^ΔPlp1^ N=3. Statistical significance determined by unpaired, two-tailed Student’s t test on animal averages. **i** Example image from pSS cortex of a Caspr^+^, CNP1^+^ paranode under MBP^+^ myelin sheath. Created in BioRender. Emery, B. (2024) BioRender.com/k49b495.

### Loss of *Fbxw7* in mature OLs results in ectopic ensheathment of neuronal cell bodies with myelin

Myelination within the CNS is highly targeted, with populations of axons displaying preferential degrees of myelination^22,34,35^ and OLs typically avoiding myelinating structures such as blood vessels and neuronal cell bodies^36–38^. There are also tight regional borders of myelination, as occurs in the cerebellum, which has distinct myelinated (nuclear layer) and non-myelinated (molecular layer) layers^39^. Exactly how this selective process is regulated is not well understood. We found that loss of *Fbxw7* in mature OLs did not alter the tight delineation of myelination between the nuclear and molecular layers of the cerebellum, with the molecular layer remaining unmyelinated (Supplemental Fig. 4a). Strikingly, however, Fbxw7^ΔPlp1^ animals displayed a significant number of granule cell bodies within the nuclear layer wrapped in MBP^+^ membrane, which drastically increased over time post-TAM (Fig. 4a-d). This mistargeting of myelin appeared to be selective to the cerebellar granule cell population; we did not observe any neuronal cell bodies wrapped in myelin in the cortex, nor did we find any other structure, like blood vessels, wrapped in MBP^+^ membrane in any regions of the CNS analyzed. Additionally, we did not observe any change in the number of OLs in our Fbxw7^ΔPlp1^ cerebellums compared to controls (Fig. 4e). It is important to note that Bergman glia in the cerebellum express *Plp1* and, therefore, may have undergone *Fbxw7* recombination in our Fbxw7^ΔPlp1^ animals^40^. While we cannot exclude *Fbxw7* KO Bergman glia as a contributing factor in our granule cell body ensheathment, we did not observe any obvious change in the number or morphology of the Purkinje cells, which are supported by Bergman glia, at 6 months post-TAM in the cerebellum (Supplemental Fig. 4a). Additionally, we found no change in GFAP expression (expressed by Bergman glia and astrocytes in the cerebellum) or reactivity of microglia (Supplemental Fig. 4b). Although the ensheathment of neuronal cell bodies appeared to be selective to the cerebellum in Fbxw7^ΔPlp1^ mice, we found that *fbxw7^vo86^* mutant zebrafish displayed a significant number of neuronal cell bodies in the spinal cord wrapped in *mbp*^+^ membrane, both in the stable *Tg(mbp-nls:EGFP); Tg(mbp:EGFP-caax)* transgenic background (Fig. 4f-h), and when OLs were mosaically labelled with a *sox10*-myrEGFP plasmid (Fig. 4i). While this phenotype may be due to the increase in OLs numbers in the zebrafish spinal cord, since it was also observed in out mouse models, it suggest the possibility that loss of FBXW7 broadly dispose OLs to mistarget their myelin to neuronal cell bodies.

**Fig. 4.**
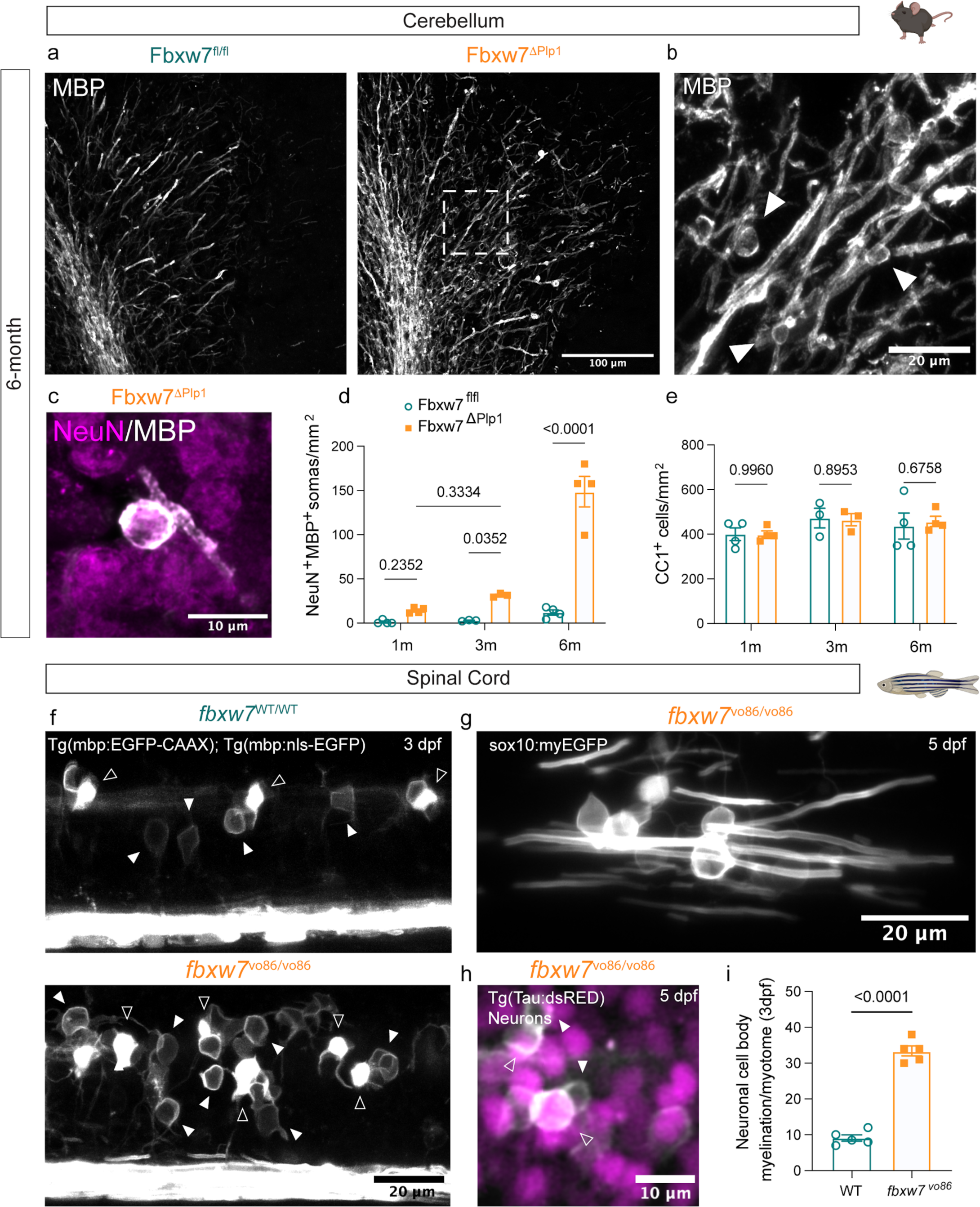
Loss of *Fbxw7* results in ectopic myelination of neuronal cell bodies in the cerebellum. **a** Representative images of anti-MBP stained cerebella from Fbxw7^fl/fl^ and Fbxw7^ΔPlp1^ animals at 6 months post-TAM. **b** High magnification image of MBP staining in the nuclear layer of a Fbxw7^ΔPlp1^ animal showing cupped myelin structures, which surround NeuN+ nuclei (shown at higher magnification in **c**). **d** Quantification of NeuN^+^ cells wrapped in MBP positive membrane in the white matter tracts and the nuclear layer of the cerebellum of Fbxw7^fl/fl^ and Fbxw7^ΔPlp1^ mice at 1, 3, and 6 months post-TAM. **e** Quantification of CC1^+^ OLs in the white matter and nuclear layer of Fbxw7^fl/fl^ and Fbxw7^ΔPlp1^ mice at 1, 3, and 6-months post-TAM. For **d** and **e**, N=4 at 1m, N=3 at 3m, N=4 at 6m. Data shown as average ± SEM. Statistical significance determined by two-way ANOVA. **f** Spinal cords of *fbxw7*^vo86^ and WT control zebrafish in *Tg(mbp:eGFP-caax)* and *Tg(mbp:nls-eGFP)* transgenic backgrounds live-imaged at 3 dpf. **g** *fbxw7^vo86/vo86^* mutant embryo injected with a plasmid driving EGFP under the *sox10* promoter at the single cell stage showing substantial wrapping of cell bodies by individual EGFP^+^ OLs at 5 dpf. **h** Representative image of an *fbxw7*^vo86^ embryo in *Tg(mbp:eGFP-caax)* and *Tg(nbt:dsRED)* (neurons) backgrounds showing neuronal cell bodies in the spinal cord wrapped in *mbp^+^*membrane. Hollow arrowheads denote OL somas, solid arrowheads denote neuronal cell body wrapping. **i** Quantification of wrapped neuronal cell bodies in the spinal cord in both *fbxw7*^vo86^ and WT controls at 3 and 5 dpf. Average ± SEM, N = 5 (larvae). Statistical significance determined by unpaired, two-tailed Student’s t test. Created in BioRender. Emery, B. (2024) BioRender.com/o45h010/k49b495.

### FBXW7 binds and degrades the N-terminus of MYRF

FBXW7 is a recognition subunit of the SKP1-Cullin-Fbox (SCF) E3 ubiquitin ligase complex. Its role is to recognize protein substrates following their phosphorylation at a phosphodegron motif, bringing them into the complex for ubiquitin tagging and subsequent proteasomal degradation^27,28,31,41^. This raised the question of which FBXW7 substrates are dysregulated in OLs after deletion of *Fbxw7* to result in accelerated and ectopic myelin formation. As a preliminary analysis, we selected a set of known FBXW7 substrates including mTOR, JNK, and cJun that are also known to regulate myelination^24,25,27,42^ and screened them by western blot in si*Fbxw7-*treated rat primary OL cultures. Surprisingly, we found no detectable changes in protein levels of mTOR, p-mTOR^ser2448^, JNK, or cJun (Supplementary Fig. 5a). To screen for FBXW7 substrates in OLs in an unbiased manner, we designed a dominant-negative FLAG-tagged version of FBXW7 missing its F-Box domain, driven under the CMV promoter (Fig. 5a). The F-Box domain of FBXW7 is required for its interaction with the SCF-E3 complex, allowing FBXW7 to disengage from its substrates^30,41^. Deletion of the F-Box domain while leaving the substrate recognition domains (WD40 repeats) intact results in a buildup of the 3xFLAG-Fbxw7^ΔF-Box^ protein bound to its substrates, allowing for effective protein-protein pulldown^30^. We electroporated primary rat OPCs with CMV-3xFLAG-Fbxw7^ΔF-Box^ or pMax-GFP controls and differentiated them for 3 days (at which point approximately 60-70% of the Olig2^+^ cells are MBP^+^). We then lysed cells, performed co-immunoprecipitation (co-IP) with an anti-FLAG antibody and assessed the eluted proteins using unbiased Liquid Chromatography-Mass Spectrometry (LC-MS). FBXW7 and MYCBP2, a known E3-independent negative regulator of FBXW7^43^, were the most highly enriched proteins by LC-MS, validating the effectiveness of our pulldown. We also found that Fbxw7^ΔF-Box^ bound MAP1B (a microtubule-associated protein), RAE1 (an RNA export protein), MYRF (a pro-myelination transcription factor), and MYO1D (an unconventional myosin; Fig. 5b). Of these targets, MYRF seemed the best-placed to mediate the phenotypes seen following loss of *Fbxw7*, and so it became our subsequent focus.

**Fig. 5.**
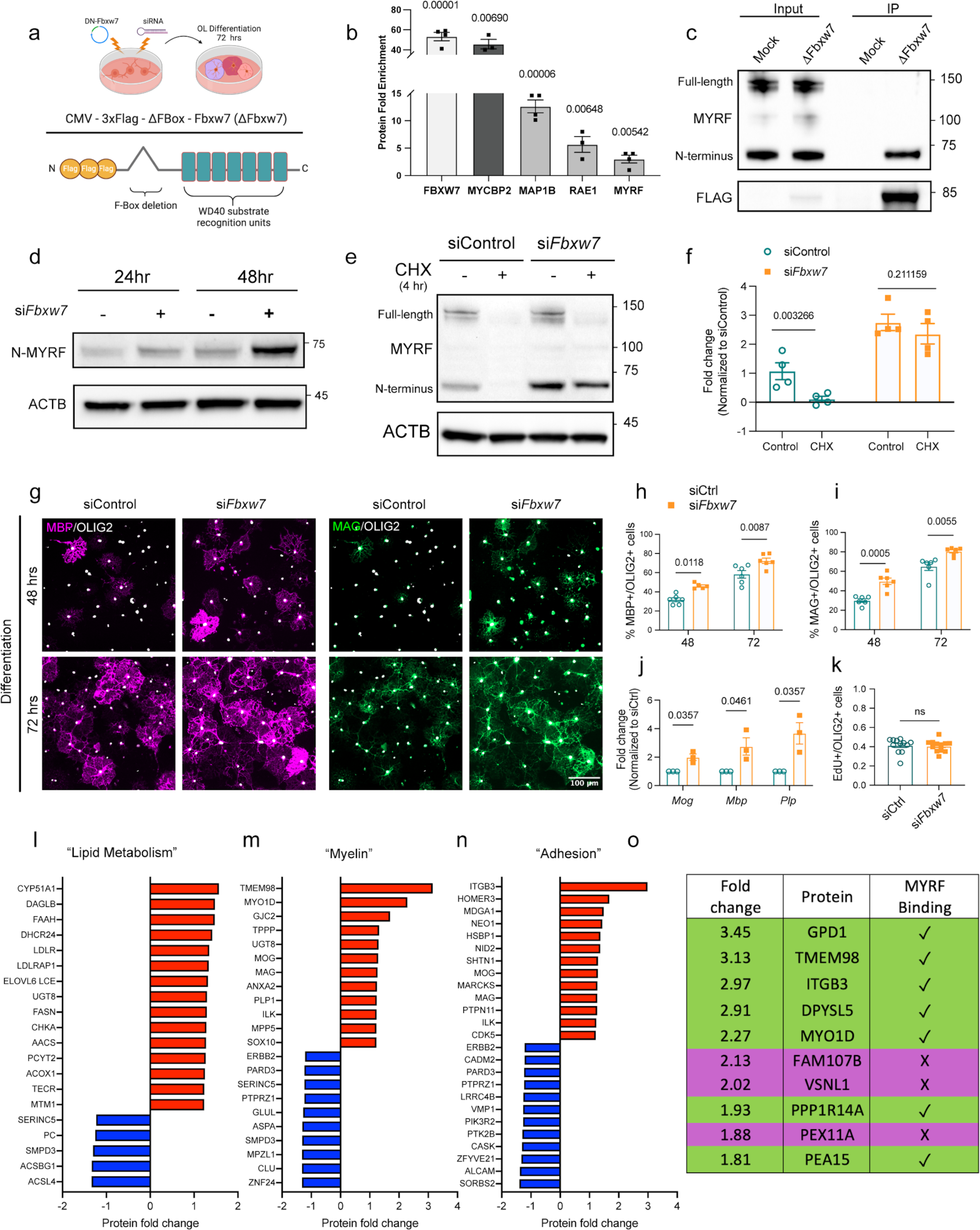
FBXW7 binds and degrades the N-terminal MYRF. **a** Schematic showing workflow of primary rat OPC isolation, expansion, and electroporation with either a dominant-negative Fbxw7 construct (CMV-3xFLAG-Fbxw7^ΔFBox^) or pooled siRNAs against *Fbxw7* or non-targeting controls. **b** LC-MS peptide counts for significant proteins enriched by anti-FLAG pull-down in 3xFlag-Fbxw7^ΔFBox^ (ΔFbxw7) electroporated cells normalized to GFP electroporated controls. Average ± SEM, N = 4 (independent cell isolations). Statistical significance determined by multiple unpaired, two-tailed Student’s t test. **c** IP-western blot for MYRF following pull down of ΔFBXW7 from cultured rat OLs at 72h differentiation. **d** Western blot analysis of the N-MYRF cleavage product in pooled siRNAs against *Fbxw7* or non-targeting controls at 24 or 48 hr differentiation. **e** Western blot of MYRF in siRNA electroporated OLs treated with cycloheximide (CHX) for 4 hours. Relative intensity of the N-MYRF band quantified in **f**. Average ± SEM, N = 4 (independent cell isolations). Statistical significance determined by unpaired, two-tailed Student’s t test. **g** Representative images of MBP and MAG expression in cultured OLs differentiated for 48 or 72 hours after electroporation with siControl or si*Fbxw7*. **h,i** MBP+ and MAG+ OLs normalized to total OLIG2+ cells. Average ± SEM, N = 3 (independent cell isolations) with 2 technical replicates (coverslips) per isolation. Statistical significance determined by two-way ANOVA. **j** qRT-PCR for myelin genes on siRNA treated cells. Average ± SEM, N = 3 (independent cell isolations) with 2 technical replicates (qRT-PCR). Statistical significance determined by multiple unpaired, two-tailed Student’s t test. **k** Quantification of EdU incorporation in primary rat OPCs following siControl or si*Fbxw7* electroporation. Average ± SEM, N = 3 (independent cell isolations) with 4 technical replicates (coverslips). Statistical significance determined by unpaired, two-tailed Student’s t test. Significant **l-n** LC-MS proteins with +/- >1.2 fold change were sorted by gene ontology (GO) terms “lipid metabolism”, “myelin” and, “adhesion”. Table of the top 10 enriched proteins by TMT-LS/MS in si*Fbxw7* electroporated OLs relative to siControl electroporated OLs at 3 days differentiation. Proteins with 1 or more MYRF ChIP-Seq binding domains within 50 KB of the transcription start site of their corresponding gene were identified based on previously published ChIP-Seq data^45^. Created in BioRender. Emery, B. (2024) BioRender.com/w60g354.

MYRF is initially produced as an endoplasmic reticulum (ER)-bound transmembrane protein. It undergoes a self-cleavage event allowing the N-terminal cleavage product (N-MYRF) to translocate to the nucleus, where it works with SOX10 at the enhancers of many essential myelin genes to promote their transcription^44–47^. MYRF levels are tightly controlled within the OL lineage, and its expression is essential for OL differentiation as well as the production and maintenance of compact myelin^13,14^. Notably, MYRF has recently been reported as an FBXW7 target in hepatocarcinoma cells, suggesting the interaction may be broadly conserved across cell types^30^.

To confirm the MYRF interaction with FBXW7 in primary OLs, we performed western blots on 3xFLAG-Fbxw7^ΔF-Box^ co-IPs. Pull-down with anti-FLAG strongly enriched for endogenous N-MYRF, but not full-length MYRF (Fig. 5c). To determine the effects of this interaction on MYRF levels, we electroporated primary rat OPCs with si*Fbxw7* and siControl, differentiated them for 24-48 hours, and blotted for endogenous N-MYRF. Knockdown of *Fbxw7* led to a substantial increase in the levels of N-MYRF, consistent with FBXW7’s role in proteasomal degradation^48^ (Fig. 5d). To confirm that the increased N-MYRF levels were due to decreased degradation in the absence of FBXW7, we treated siControl and si*Fbxw7* OL cultures with cycloheximide (CHX) to inhibit protein translation. We found that after 4 hours of CHX treatment, control cells had degraded the majority of both full-length MYRF and N-MYRF. In contrast, *Fbxw7* knockdown cells showed little reduction in N-MYRF levels, but near complete loss of the full-length protein (presumably due to clearance via self-cleavage) (Fig. 5e, f). Together, these findings strongly supported a role for FBXW7 in N-MYRF degradation.

When analyzing the N-MYRF blots we noticed two distinct molecular weights of N-MYRF separated by ∼2 kDa, with the higher molecular weight band becoming more prevalent with *Fbxw7* knockdown (Fig. 5d, e). Nakayama and colleagues found that phosphorylation of MYRF at serine 138 and 142 by GSK3β was required for FBXW7 to interact with N-MYRF^30^. To determine whether build-up of a phosphorylated form accounted for the observed change in molecular weight, control and si*Fbxw7* OL lysates were treated with a lambda phosphatase, and molecular weights were evaluated by western blot (Supplementary Fig. 5b). We found that treatment with phosphatase resulted in a near total loss of the larger molecular weight N-MYRF in both control and si*Fbxw7* OLs, consistent with phosphorylated N-MYRF constituting the majority of the increased N-MYRF in our *Fbxw7* knockdown OLs. To determine if GSK3β is the kinase responsible for phosphorylating the phospodegron motif in N-MYRF to induce its interaction with FBXW7, as shown in hepatocarcinoma cell lines^30^, we electroporated OPC cultures with pooled siRNAs against *Gsk3b* and differentiated them for 3 days. Although GSK3β protein was robustly downregulated, we found no change in the level of N-MYRF or corresponding myelin proteins (Supplemental Fig. 5c). Together, these data suggest that although N-MYRF is phosphorylated in OLs and that this phosphorylated form is the target of FBXW7, GSK3β is not the primary kinase responsible for targeting MYRF for FBXW7-mediated degradation in OLs.

Given MYRF’s well defined role in OL differentiation and myelination we next wanted to investigate the functional consequences of elevated MYRF levels in *Fbxw7* knockdown OLs. si*Fbxw7*-electroporated OLs differentiated for 48-72 hours showed a significant increase in the proportion of MBP^+^ and myelin-associated glycoprotein (MAG)^+^ cells compared to controls (Fig. 5g-i). In addition to an increased proportion of cells expressing myelin proteins, cultures also showed significant increases in *Mbp*, *Mag*, and *Plp1* mRNA as assessed by qRT-PCR (Fig. 5j). To determine if loss of *Fbxw7* in OPCs was sufficient to induce differentiation in the presence of PDGF-AA, si*Fbxw7* and control OPCs were kept in proliferation media for 2 days after siRNA electroporation to allow for effective knockdown, then pulsed with 5-ethynyl-2’-deoxyuridine (EdU) for 6 hours. Within that time, approximately 40% of OPCs had undergone a round of division, with no significant change seen in EdU incorporation between si*Fbxw7* or siControl treated OPCs (Fig. 5k), indicating that loss of *Fbxw7* in OPCs was not sufficient to induce *Myrf* expression and the transition to a post-mitotic OL. These results indicate that once OPCs begin to differentiate and express *Myrf*, FBXW7 serves to regulate N-MYRF protein levels to control the balance and timing of OL myelination.

To further understand the consequences of loss of *Fbxw7* on the OL proteome, we performed LC-MS on lysates from si*Fbxw7* and siContol electroporated rat OLs at 3 days of differentiation. Cell lysates were labeled with tandem mass tags (TMT), pooled, and run through LC-MS. Over 2,700 proteins were sequenced with an R^2^ value of 0.99-1 within treatment groups. Select enriched proteins with validated antibodies were confirmed by western blot (Supplemental Fig. 5d). We found that *Fbxw7* knockdown in primary OLs resulted in significant changes in 253 proteins with a false discovery rate (FDR) <0.01 and 426 proteins with an FDR <0.05. Within the 253 proteins with an FDR <0.01, 158 proteins showed increased levels with *Fbxw7* knockdown and 95 showed reduced levels. Proteins with significant changes and a fold change greater than ±1.2 were sorted by Gene Ontology functions “lipid metabolism,” “myelin,” and “adhesion” (Fig. 5l-n) to provide a list of proteins with potential roles in FBXW7-dependent myelination.

During OL differentiation, N-MYRF directly binds the enhancer regions of genes underpinning myelination, with enrichment of N-MYRF chromatin-immunoprecipitation (ChIP) peaks seen within 50 kb of the transcription start sites of genes induced during OL differentiation^45^. Notably, when the proteins were ranked by fold change following *Fbxw7* knockdown, 7 of the top 10 upregulated proteins and 73% of all proteins with fold change >1.5 had a predicted N-MYRF binding motif within 50 kb of the transcriptional start site of their corresponding gene (Fig. 5o). When we assessed RNA from corresponding samples by qPCR, we found that all the top 10 enriched proteins showed significant increases in their transcript levels with *Fbxw7* knockdown, suggesting that many of the changes in protein abundance following loss of *Fbxw7* were secondary to elevated MYRF transcriptional activity (Supplemental Fig. 5e).

### Loss of *Fbxw7* in OLs increases nuclear MYRF levels *in vivo*

We next sought to determine whether the myelin changes seen in Fbxw7^ΔPlp1^ mice may be mediated by elevated MYRF levels. Since the antibody we used to detect MYRF recognizes an epitope within the N-terminal 100 amino acids of MYRF, it recognizes both the full-length form, which is bound to the ER, and the N-terminal cleavage product, which is translocated to the nucleus^45,46^. To determine the abundance of each, we use the cellular localization of the cytoplasmic (full-length) or nuclear (N-terminal) for quantification of MYRF levels in Fbxw7^ΔPlp1^ and control optic nerve OLs. In Fbxw7^ΔPlp1^ animals, we found a significant increase in the levels of nuclear-MYRF relative to their control littermates at 1- and 3-months post *Fbxw7* deletion (Fig. 6a-c). We observed similar increases in the intensity of nuclear MYRF staining in the cerebellum and corpus callosum of Fbxw7^ΔPlp1^ mice (Supplemental Fig. 6). Interestingly, the levels of nuclear MYRF in Fbxw7^ΔPlp1^ optic nerves returned to control levels by 6 months post-TAM. This was associated with a significant reduction in the ratio of cytoplasmic to total (nuclear + cytoplasmic) MYRF levels (Fig. 6d). We believe this reduced ratio of full-length MYRF to nuclear MYRF may represent a homeostatic response by the OL to control MYRF protein levels in the absence of FBXW7 by down-regulating *Myrf* transcription.

**Fig. 6.**
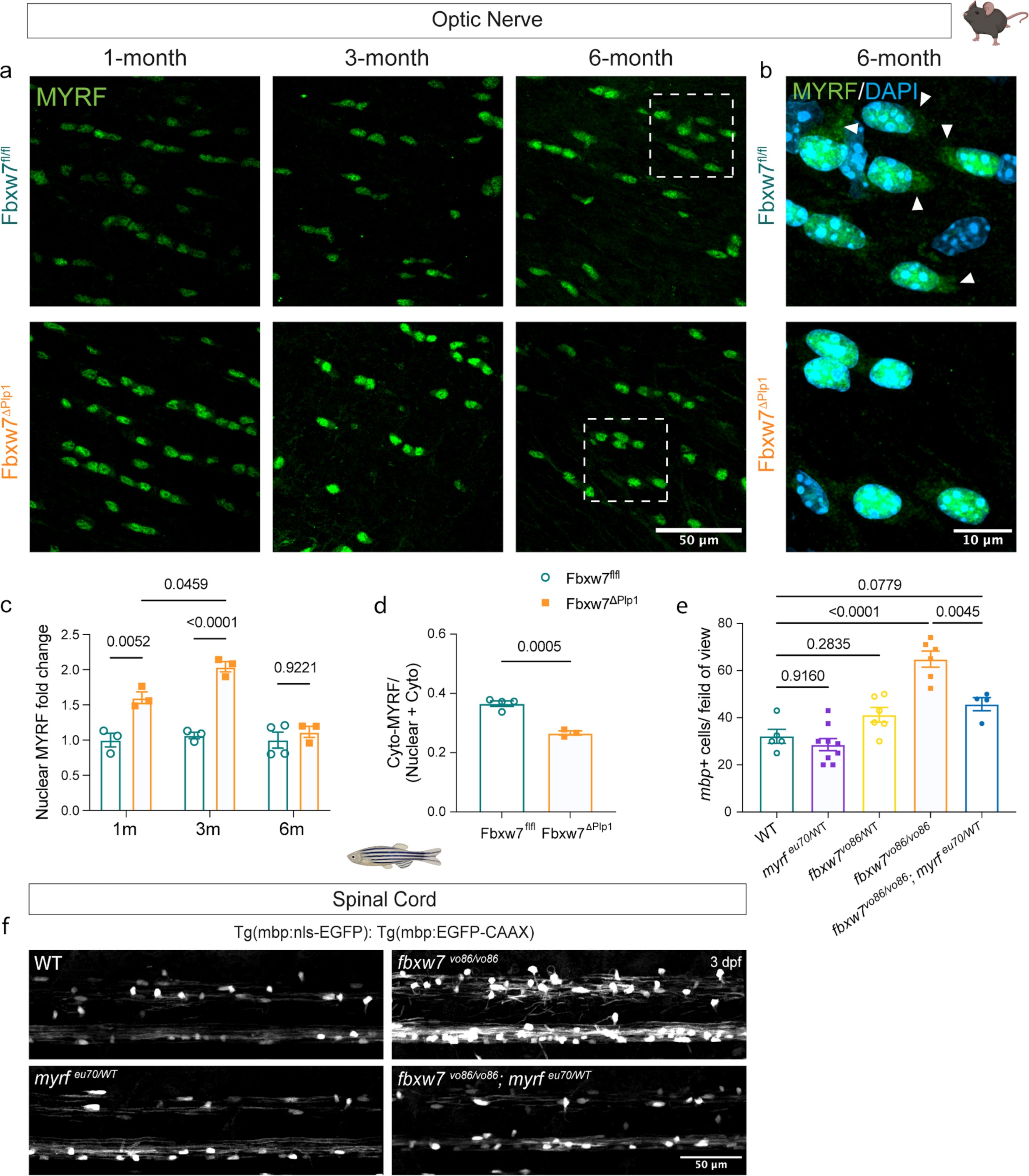
FBXW7 regulates OL MYRF levels *in vivo*. **a** Representative images of optic nerves from Fbxw7^fl/fl^ and Fbxw7^ΔPlp1^ mice at 1-, 3-, and 6-months post-TAM stained for MYRF. **b** High resolution images showing cytoplasmic localization of MYRF in the optic nerve OLs of 6 months post-TAM animals. Arrowheads indicate cytoplasmic localization of MYRF (likely uncleaved precursor) in the Fbxw7^fl/fl^ control. **c** Quantification of nuclear MYRF intensity in Fbxw7^ΔPlp1^ nerves normalized to controls at 1-, 3-, and 6-months post-TAM. For both genotypes, N=4 at 1m, N=3 at 3m, N=4 at 6m. Data shown as average ± SEM. Statistical significance determined by two-way ANOVA. **d** Ratio of cytoplasmic relative to total (cytoplasmic and nuclear) MYRF at 6 months post-TAM. Statistical significance determined by unpaired, two-tailed Student’s t test. **e** Quantification of *mbp:nls^+^* OLs in the zebrafish dorsal spinal cord of each genotype. WT (N=5), *myrf^eu70/^*^WT^ (N=9), *fbxw7^vo86^*^/WT^ (N=6), *fbxw7^vo86/vo86^*(N=6), *fbxw7^vo86/vo86^; myrf^eu70/^*^WT^ (N=4). Average ± SEM, statistical significance determined by two-way ANOVA. **f** Representative images of the spinal cords of *fbxw7^vo86/vo86^and fbxw7^vo86/vo86^; myrf^eu70/^*^WT^ zebrafish on *Tg(mbp:eGFP-caax)* and *Tg(mbp:nls-eGFP)* transgenic backgrounds at 3 dpf. Created in BioRender. Emery, B. (2024) BioRender.com/o45h010/k49b495.

The elevation of N-MYRF levels in Fbxw7^ΔPlp1^ mutants, combined with the elevated mRNA and protein levels of many known MYRF targets in si*Fbxw7* electroporated OLs in culture, strongly suggested that elevated MYRF levels may underlie the precocious myelination seen following loss of *Fbxw7.* To test if *myrf* is epistatic to *fbxw7* in OLs, we crossed the *fbxw7^vo86/vo86^* line to a *myrf* mutant line (*myrf^eu70^*^/WT^)^49^ to determine if reducing *myrf* levels could suppress *fbxw7* mutant phenotypes *in vivo*. At 3 dpf, we observed a significant reduction in the number of *mbp:nls-EGFP^+^* OLs in the spinal cord of *fbxw7^vo86/vo86^, myrf^eu70^*^/WT^ animals compared to *fbxw7^vo86/vo86^* mutants alone (Fig. 6e, f). These data, along with our work in primary OL cultures and conditional KO mouse models show that FBXW7 negatively regulates N-MYRF in OLs to control many facets of OL biology, from OL sheath length, paranodal organization, to long-term homeostatic maintenance of myelin.

## Discussion

### FBXW7 regulation of myelin homeostasis

Here we present evidence that once cells are committed to the OL lineage, FBXW7 regulates OL myelin capacity, organization, and homeostasis, in part through the negative regulation of the N-MYRF transcription factor. With the temporal control of inducible conditional knock-out mice we found that loss of *Fbxw7* in myelinating OLs resulted in increased myelin sheath length, severe myelin outfolds, disorganized paranodes, and surprisingly, wrapping of neuronal cell bodies in the cerebellum. These phenotypes are particularly striking since *Fbxw7* was targeted in mature OLs, indicating that inactivation of FBXW7 reinitiates aspects of myelin growth in the adult CNS. While myelin sheath plasticity has been reported in the context of axonal activity^32,50,51^, the underlying mechanisms that regulate these changes have not been fully characterized. Because loss of *Fbxw7* results in changes to many adhesion and cytoskeleton proteins, it is possible that FBXW7-mediated negative regulation of OL proteins may represent one of these underlying mechanisms of plasticity.

Ensheathment of neuronal cell bodies occurs with global deletion of the inhibitory myelin-guidance protein JAM2^38^ and also occurs in zebrafish when OL numbers exceed their normal balance to receptive axons^52^. It is entirely possible that the increase in ensheathed neuronal cell bodies we observe in the *fbxw7^vo86^* zebrafish is a consequence of increased OPC specification and OL numbers^23^. In contrast, myelin ensheathment of cerebellar granule cell bodies of Fbxw7^ΔiPlp1^ mice was not accompanied by an increase in the density of OLs, so is unlikely to be mediated by a mismatch between the myelinating cells and their targets. Interestingly, we did not observe any wrapping of cell bodies or other structures in the cerebral cortex of Fbxw7^ΔPlp1^ animals. Why cerebellar granule cell bodies were the only observed neurons ensheathed by myelin in the *Fbxw7* conditional knockouts is puzzling. It is possible the granular cell body ensheathment is secondary to myelin outfolding, as reported previously following N-WASP inactivation within the OL lineage^53^. In this scenario, perhaps the density and/or size of cerebellar granule cells makes them susceptible to ensheathment by redundant myelin outfolds. Indeed, ensheathment of granule cell bodies has also been observed in normal development of the toad cerebellum^54^. Alternatively, loss of *Fbxw7* in OLs could lead to over- or under-expression of targeting molecules that would normally prevent myelination of granule cells (see below).

In Schwann cells, FBXW7 regulates myelin sheath thickness, with no obvious change to sheath length^25^. In OLs these roles seem to be reversed, with FBXW7 regulating myelin sheath length but not thickness. Although this may be due to inherent differences in the biology of these two cell types, it may also be due to the differences in the tools used to evaluate its function. In our prior Schwann cell work, *Fbxw7* was constitutively deleted in development using Desert hedgehog Cre (Dhh-Cre), which is expressed in Schwann cell precursors as early as E12.5^25,55^. In contrast, here we used a TAM-inducible system to delete *Fbxw7* from myelinating OLs in 8-week-old Plp1-CreERT mice. Whether the effects of *Fbxw7* on myelination would change depending on the timing of OL deletion remains unclear, but is an exciting proposition for future work.

### Divergent FBXW7 targets across the OL lineage

Our studies and others highlight the complicated role FBXW7 plays in myelinating cell biology across species and cell types. *Fbxw7* is widely expressed in most, if not all cell types in the CNS, and its biological functions depend on the available substrates within each cell type^41,48,56^. Therefore, FBXW7 is likely to have distinct targets and diverging roles at different stages of the OL lifespan. For instance, FBXW7 negatively regulates NOTCH levels in neural precursor cells (NPCs), to control OPC specification in zebrafish^23^. The same group also found that in later stages of the OL lineage, FBXW7 negatively regulates mTOR to control myelination in the spinal cord of zebrafish. Likewise, we previously showed that in the mammalian PNS, FBXW7 regulates early Schwann cell numbers, axonal ensheathment, and myelin thickness in an mTOR-dependent manner^25^. In contrast, here we present evidence that in mammalian OLs, mTOR was not a direct target of FBXW7, with knockdown of *Fbxw7* resulting in no detectable change in mTOR protein abundance, phosphorylation, or downstream signaling. Using an unbiased pull-down and LC-MS approach, we instead identified several direct FBXW7-interacting proteins in primary OLs; MYCBP2, MAP1B, RAE1, and MYRF. Of these, MYCBP2 was a previously identified FBXW7 interactor that likely inhibits its activity independent of its E3 complex^43^. Although we focus here on MYRF, MAP1B and RAE1 also represent intriguing FBXW7 targets within myelinating cells.

### Regulation of MYRF

As a critical regulator of OL differentiation and myelination, MYRF levels appear to be tightly regulated within the OL lineage. Not only is it subject to tight transcriptional regulation by SOX10 and ZFP24^57^, but its mRNA is subject to negative regulation by miR-145-5p in OPCs, presumably to discourage premature differentiation^58^. In contrast, the posttranslational mechanisms regulating N-MYRF activity remain incompletely understood. The full length MYRF protein trimerizes and self-cleaves to enable release of the active transcription factor^45,46,59^. Although this process is negatively regulated by TMEM98, the protein product of one of MYRF’s own target genes^59,60^, the degree to which TMEM98 negatively regulates the production of the N-MYRF transcription factor at endogenous levels remains unclear. Our findings that FBXW7 negatively regulates N-MYRF levels in primary OLs corroborate the finding that FBXW7 targets N-MYRF in mHepa cells^30^, suggesting a conserved regulatory mechanism. Indeed, the fact that OL numbers could be normalized in the *fbxw7^vo86^* fish by *myrf* haploinsufficiency highlights N-MYRF as a central FBXW7 target within the OL lineage. In contrast to mHepa cells, however, in primary OLs, the FBXW7 interaction does not seem to depend on the phosphorylation of the phospho-degron motif by GSK3b. The intracellular pathways that initiate N-MYRF turnover by FBXW7 will be important to determine in future work.

Notably, the dysregulated proteins seen in *Fbxw7* knockdown OLs included many cell adhesion and cell surface proteins. Previous research has highlighted the connection between adhesion molecules and proper myelination. For instance, Djannatian et al. found that when contactin-associated protein (*Caspr*), contactin-1 (*Cntn1*), neurofascin (*Nfasc155*), and myelin-associated glycoprotein (*Mag*) were globally deleted in both the zebrafish and mouse CNS, similar myelin phenotypes were observed as in Fbxw7^ΔPlp1^ animals^61^. They found that the loss of these adhesion proteins resulted in outfolds, double myelinated axons, and loss of paranodal loop organization. Additionally, they also observed neuronal cell body myelination in the zebrafish spinal cord. It is exciting that within our model, where *Fbxw7* deletion is restricted to myelinating OLs, we see such similar phenotypes when compared to global disruption of adhesion proteins in the CNS.

In summary, we have shown that FBXW7 is an evolutionary conserved regulator of OL myelin capacity and homeostasis. We found that FBXW7 regulates myelination by controlling sheath elongation independent of myelin wraps and is required for long-term maintenance of myelin integrity and paranodal organization within the adult CNS, in part, through its negative regulation of N-MYRF.

## Methods

### Zebrafish husbandry

All zebrafish experiments were done in compliance with the institutional ethical regulations for animal testing and research at Oregon Health & Science University (OHSU). *fbxw7^stl64^*, *fbxw7^vo86^*, and *myrf^eu70^* zebrafish were maintained as heterozygotes. Experimental larvae were generated by incrosses to yield wild-type, heterozygous, and homozygous zebrafish. To create *fbxw7^vo86/vo86^; myrf^eu70^*^/WT^ animals, *fbxw7^stl6/^*^WT^; *myrf^eu70^*^/WT^ zebrafish were outcrossed to *fbxw7^stl86/^*^+^. Zebrafish larvae are fed a diet of rotifers and dry food (Gemma 75) from 5 days post-fertilization (dpf) until 21 dpf. From 21 dpf until 3 months, fish are fed using rotifers and dry food (Gemma 150). Adult fish are maintained and fed with brine shrimp and dry food (Gemma 300). For larval zebrafish studies, sex cannot be considered as a biological variable as sex has not yet been determined.

|  |
| --- |
| Zebrafish transgenic lines |
| *Tg(mbp:eGFP-caax)*<sup>62</sup> |
| *Tg(olig2:dsRED)*<sup>63</sup> |
| *Tg(mbp-nls:eGFP)*<sup>64</sup> |
| *Tg(nbt:dsRED)*<sup>65</sup> |

### Generation of *fbxw7^vo86^* zebrafish mutants

CRISPR-Cas9 was used to generate genetic mutants in zebrafish. The CHOPCHOP web tool^66^ was used to select target sites, and individual sgRNAs were synthesized using MEGAshortscript T7 Transcription kit (Thermo Fisher). The sgRNA GATGTAATCCGTCGTCTCTGTGG was mixed with Cas9 Nuclease (Integrated DNA Technologies) to a final concentration of 50 ng/mL sgRNA and 1 mg/mL of Cas9 protein and injected into one-celled zygotes at a volume of 1-2 nL. Progeny of injected F0 generation were screened for the presence of inherited indels resulting in frameshifts or truncations by PCR, and these F1 progenies were used to establish stable mutant lines. Genotyping for both larval and adult zebrafish was performed by digesting tissue in tris buffer with proteinase-K overnight at 55°C. PCR was performed with GoTaq DNA Polymerase (Promega, M300A). For *fbxw7^vo86^* genotyping, PCR with (F-AAAATAGGGGCTTGCTCTGG, R-AAGTCCAGTTAAATTGAGAAGCC) was used to amply a 530 bp regain around indels. PCR products were digested with 10U of BsmBI-v2 (NEB, R0580) at 55°C overnight and resolved on a 2% agarose gel.

### Mosaic labeling and cell-type specific CRISPR-Cas9 gene disruption in zebrafish oligodendrocytes

For mosaic labeling of oligodendrocytes (OLs), *fbxw7^vo86/^*^+^ zebrafish were incrossed, and fertilized one-cell zygotes were injected with 1-2 nl of a solution containing 10 ng of sox10:EGFP-caax plasmid, 25 ng of Tol2 transposase mRNA, 0.02% phenol red and 0.2 M KCl. Embryos were genotyped after imaging as described above. For cell-type specific CRISPR-Cas9 mediated gene disruption we utilized methods as previously described^32^. Briefly, sgRNAs targeting *fbxw7* exon 5 (GATGTAATCCGTCGTCTCTGTGG) and exon 7 (GCTGCCTGAAGCAGATCCTTTGG) were cloned into 10xUAS:myrmScarlet-p2A-Cas9, U6:sgRNA1;U6:sgRNA2 backbones and injected into *Tg(sox10:Kalta4)*^67^ fertilized embryos at the one-cell stage. Empty backbones were used as controls. At desired timepoints, fish were anesthetized with 600 μM tricaine (TRS5, Pentair), screened for fluorescence, embedded laterally in 1.5% low-melting-point agarose (A9414, Sigma), and imaged with a 20x dipping objective on a ZEISS LSM 980 with Airyscan 2. Sheaths were analyzed using ImageJ.

### Mouse husbandry and tamoxifen (TAM) administration

All mice were housed in OHSU animal facilities, maintained in a pathogen-free temperature and humidity-controlled environment on a 12-hour light/dark cycle. All procedures were approved by the OHSU Institutional Animal Care and Use Committee. Fbxw7^fl/fl^ mice were purchased from Jackson Laboratories (B6;129-*Fbxw7^tm1Iaai^*/J, JAX: 017563) and crossed to Plp1-CreERT mice (B6.Cg-Tg[Plp1-cre/ERT]3Pop/J, JAX:005975). CreERT negative littermates served as controls. Genotypes were determined by PCR analysis using established primers for each line and were revalidated at experimental endpoints. All experiments were conducted in both male and female mice. For TAM injection, 8-week-old mice were dosed with 100mg/kg tamoxifen (Sigma T5648, dissolved at 20 mg/ml in corn oil) for five consecutive days via intraperitoneal injection.

### Tissue processing

Mice were terminally anaesthetized with ketamine (400 mg/kg) and xylazine (60 mg/kg) before being transcardially perfused with 20 mL of phosphate buffered saline (PBS) and 40 mL of 4% paraformaldehyde (19210, Electron Microscopy Sciences) in PBS. For immunofluorescence (IF), tissues were post-fixed in 4% paraformaldehyde in PBS (2 hours for optic nerves, overnight for brains) and cryopreserved in 30% sucrose for at least 72 hours at 4°C. Cortical flat mounts were processed as previously described^35^. Cryopreserved tissue was embedded in OCT (4583, Sakura), frozen on dry ice and stored at -80°C until sectioning on a cryostat (Leica CM3050-S). Cryosections (12 μm thickness for brain, 16 µm for optic nerve) were mounted on Superfrost Plus slides (1255015, Fisher Scientific) and stored at -80°C. Tissue for electron microscopy was post-fixed in 2% paraformaldehyde (15710, Electron Microscopy Sciences) with 2% glutaraldehyde (16310, Electron Microscopy Sciences).

### Immunofluorescence

Slides stored in -80°C were air dried for at least 2 hours before being rehydrated in 1x PBS. For MBP staining, tissue was delipidated by treating slides with ascending and descending ethanol solutions (50%, 75%, 85%, 95%, 100%) before being washed 3x in 1x PBS. Slides were blocked for 1 hr at room temperature with 10% fetal calf serum (SH30910.03, Cytiva) with 0.2% Triton X-100 (10789704001, Sigma). Primary antibodies were applied overnight in 1x PBS, 5% fetal calf serum and 0.2% Triton X-100 in a sealed container containing water at room temperature. The following primary antibodies were used: chicken anti-MBP (1:500; MBP, Aves), mouse anti-CC1 monoclonal (1:500; OP80, Millipore), goat anti-PDGFRα (1:500; AF1062, R&D Systems), rabbit anti-Iba1 (1:1000; 019-19741, Wako), rabbit anti-GFAP (1:1000; Z0334, Dako), rabbit anti-MYRF (1:500; A16355, ABclonal), mouse anti-Calbindin1 (1:500; C9848, Sigma). Following incubation with primary antibodies, slides were washed 3x in 1x 0.2% Triton X-100 PBS before appropriate Alexa Fluor 488, 555 or 647 secondary antibodies (1:1,000; Invitrogen) were applied for one hours at room temperature. Slides were then again washed 3x with 1x 0.2% Triton X-100 PBS, washed in Milli-Q H_2_O, air dried, then coverslipped with Fluoromount G (0100-01, Southern Biotech) and imaged.

Primary rat OLs were cultured on glass coverslips, fixed for 8 minutes in 4% PFA in PBS and stained in 24-well plates as described above. We used the following primary antibodies: chicken anti-MBP (1:500; MBP, Aves), rabbit anti-OLIG2 (1:500; AB9610, Millipore), and mouse anti-MAG (1:500; AB9610, Millipore). Coverslips were mounted with ProLong Diamond (P36965, Thermo Fisher) on Superfrost Plus slides.

Cortical flatmounts were sectioned at 40 μm and stored in PBS with 0.02% NaN_3_ (sodium azide) at 4°C. For staining, tissue was blocked in 10% fetal calf serum with 0.2% Triton X-100 for 2 hours with agitation at room temperature. Flatmounts were then incubated with primary antibodies at room temperature with agitation in 1x PBS 0.2% Triton X-100 for 4 days. We used the following primary antibodies: chicken anti-MBP (1:200; MBP, Aves), rabbit anti-CASPR (1:500; 34151-001, Abcam), and mouse anti-CNPase1 (1:500; MAB326, Millipore). Tissue was washed 3x 20 min 1x 0.2% Triton X-100 PBS. Alexa Fluor 488, 555 and 647 secondary antibodies (1:1,000; Invitrogen) were applied for two days at 4°C protected from light. Tissue was washed 3x 20 min 1x 0.2% Triton X-100 PBS, slide mounted, rinsed in water, air dried, and coverslipped with Fluoromount G (0100-01, Southern Biotech). For quantification of sheath length from layer-I pSS a 280 μm x 280 μm x 30 μm images were taken of the pSS and 2 random ROIs were generated. All sheaths that passed through the ROIs were measured using ImageJ NeuroTracer in 3D.

Immunofluorescence from tissue was acquired on a ZEISS LSM 980 with Airyscan 2. Immunofluorescence from cultured OLs were imaged on Zeiss ApoTome2 at 20x. All cell counts and fluorescence intensities were quantified using ImageJ.

### Isolation, expansion, and electroporation of primary rat OPCs/OLs

Rat OPCs were isolated from P6-8 Sprague Dawley rat pups as previously described^68^. OPCs from each animal were expanded in 3x 175cm^2^ flasks for 3-4 days in the presence of 10ng/mL platelet derived growth factor-aa (PDGFAA, Peprotech 100-13A). Cells were harvested fresh for each round of experiments and used at the time of first passage. OPCs were electroporated with Amaxa Basic Nucleofector Kit for Primary Mammalian Glial Cells (VPI-1006, Lonza) with 20 nM siRNAs/5 million OPCs or 4 μg of plasmid/5 million OPCs. siRNA pools for rat *Fbxw7* (L-115782-00-0005, Horizon) and GSK3β (L-080108-02-0005, Horizon) or non-targeting controls (D-001810-10-05, Horizon) were used. Primary rat OLs were plated at 20k/coverslip for staining, 250k cells/well of a 6-well plate for RNA, and 1 million cells/60x15 mm plate for protein isolation.

### EdU incorporation and cycloheximide (CHX) treatment

Primary rat OPCs were expanded and electroporated with siRNAs and replated into proliferation media containing PDGFAA for 48 hours. Cells were pulsed with 10 μM 5-ethynyl-2’-deoxyuridine (EdU) for 6 hours. Cells were fixed with 4% PFA in PBS for 8 minutes at room temperature. Cells were stained with Click-iT EdU Cell Proliferation Kit with Alexa Fluor 647 dye (C10340, Thermo Fisher). siRNA electroporated OLs were differentiated for 3 days and treated with cycloheximide (CHX; 239763-M, Sigma) at 100 μg/mL for 4 hours to stop protein translation. Cell lysates were processed for western blot analyses as described below.

### Transmission electron microscopy

Following post-fixation in 2% paraformaldehyde (15710, Electron Microscopy Sciences) with 2% glutaraldehyde (16310, Electron Microscopy Sciences), optic nerves were stored in a buffer of 1.5% paraformaldehyde, 1.5% glutaraldehyde, 50 mM sucrose, 22.5 mM CaCl_2_ 2H_2_O in 0.1M cacodylate buffer for at least seven days. Tissue was then infiltrated with 2% osmium tetroxide (19190, Electron Microscopy Sciences) using a Biowave Pro+ microwave (Ted Pella) before dehydration in acetone and embedding in Embed 812 (14120, Electron Microscopy Sciences). 0.4 µm sections were cut on an ultramicrotome and stained with 1% Toluidine Blue (T161-25, Fisher Scientific) with 2% sodium borate (21130, Electron Microscopy Sciences). 60 nm sections were mounted on copper grids (FCF100-Cu-50, Electron Microscopy Sciences) and counterstained with UranyLess for 5 minutes followed by 3% lead citrate (22409, 22410, Electron Microscopy Sciences) for 5 minutes. Grids were imaged at 4800x on an FEI Tecnai T12 transmission electron microscope with a 16 Mpx camera (Advanced Microscopy Techniques Corp). For g-ratio analysis, 5-8 images per animal were used. Outer myelin and axon diameters for g-ratio analyses were manually traced using ImageJ.

### Cloning of dominant-negative 3xFLAG-Fbxw7^ΔF-Box^

*Fbxw7* coding sequence missing the F-Box domain with upstream 3x FLAG tags (3xFLAG-Fbxw7^ΔF-Box^) was purchased as double stranded gBlocks gene fragment from Integrated DNA Technologies (IDT) with KpnI + HindIII restriction enzyme overhangs. pCMV SPORT backbone and 3xFLAG-Fbxw7^ΔF-Box^ inserts were digested with KpnI + HindIII restriction enzymes, gel purified, and ligated with T4 DNA ligase (EL0011, Thermo Fisher). Constructs were transformed into DH5⍺ one-shot competent cells (12297016, Thermo Fisher) and purified with a PureLink HiPure plasmid maxiprep kit (K0491, ThermoFisher). pmaxGFP vector (Lonza) was used as a control for co-IP experiments.

### Immunoprecipitation of Fbxw7 dominant-negative constructs in cultured OLs

4 μg of CMV-3xFLAG-Fbxw7^ΔF-Box^ and pmaxGFP were electroporated into rat OPCs. Cells were differentiated for 3 days and then lysed with cell lysis buffer (20mM Tris pH7.5, 150mM NaCl, 1% Triton X-100, 1mM EDTA, 1mM EGTA) with cOmplete Mini Protease Inhibitor Cocktail (11836153001, Millipore) for 30 min at 4°C with rotation. Lysates were spun at 4°C for 10 min at 10,000 RPM. 5% of lysates were frozen for input controls. 4 μg of mouse anti-FLAG M2 antibody (F3165, Millipore) was added to lysates and rotated at 4°C for 2 hours. 40 μL Dynabeads Protein G (10003D, Thermo Fisher) was added to lysates and rotated for 1 hour at 4°C. Beads were sorted out with a magnetic rack and washed x5 with cell lysis buffer with rotation at 4°C for 5 mins. Proteins were released from Dynabeads Protein G beads for LC-MS by boiling in 1% SDS. Proteins for western blots were boiled with 1x Laemmli buffer.

### Western blots

For western blots on cultured OLs, plates were washed 3x with cold DPBS and lysed with RIPA buffer (50mM Tris-HCL pH 8.0, 150 mM NaCl, 1% NP-40, 0.5% Sodium deoxycholate, 0.1% SDS, 1 mM EDTA, 0.5 mM EGTA) with complete protease inhibitors (11836153001, Roche), and phosphatase inhibitors (04906837001, Roche) before being spun at 13,000g at 4°C. Protein lysate was removed and frozen at -80°C. Lysates were boiled at 98 °C in 1x Laemmli buffer for 5 min and run on Bis-Tris-gel (NP0335BOX, Invitrogen). To transfer proteins to PVDF membranes (IPVH00010, Thermo Scientific), transfer cassettes were assembled (A25977, Thermo Fisher Scientific) and filled with transfer buffer (NP0006-01, Thermo Scientific) containing 10% methanol and run at 20V for 1hr. Following transfer, blots were rinsed in 1x TBS with 0.1% Tween-20 (TBST) before blocking in 1x TBST with 5% milk powder for one hour at room temperature. Blots were probed with antibodies against MYRF (16355, ABClonal), mTOR (2983, Cell Signaling), Phospho-mTOR (Ser2448; 5536, Cell Signaling), cJun (9261L, Cell Systems), TMEM98 (14731-1-AP, Proteintech), MYO1D (ab70204, abcam), GSK3β (ab32391, abcam), DPYSL5 (CRMP5; ab36203, abcam). All antibodies for western blots were used at a concentration of 1:1000. Blots were incubated in primary antibodies diluted in 2.5% BSA (BP9706-100, Fisher Scientific) with 1% NaN_3_ in TBST overnight at 4°C. After overnight incubation, blots were washed in 3x TBST and incubated with appropriate HRP-conjugated secondary (Goat anti-rat 7077, Cell Signaling; Goat anti-mouse 7076, Cell Signaling, Goat anti-rabbit 7074, Cell Signaling) at 1:5,000 for two hours with 2.5% milk powder in TBST. Immunoreactivity was visualized using chemiluminescence (34080, Thermo Fisher Scientific) and imaged on a Syngene GBox iChemiXT. Blots were then re-probed with β-actin-HRP (1:5000; A3854, Sigma). Densitometric analysis was performed in ImageJ by quantifying the intensity of bands relative to the ACTB loading control and then normalized to background.

### Mass spectrometry and analysis

OLs electroporated with pooled siRNAs targeting *Fbxw7* or non-targeting controls were differentiated for 72 hours and lysed with eFASP buffer (4% SDS, 0.2% DCA, 100mM TEAB), frozen at -80 °C, and submitted to the OHSU proteomic core. Samples were then sonicated using Bioruptor Pico (30s on 30s off, 10 cycles), heated to 90°C for 10 mins, cooled, centrifuged, and protein concentration was determined by BCA. 55μg of protein/sample were digested with eFASP and measured by peptide assay. 18 μg peptides/sample were labeled with TMT 11-plex, normalized, and pooled. Pooled TMT samples were then run on an 18 fraction 2dRPRP LC/MS on Orbitrap Fusion. Data was then analyzed with COMET/PAWS pipeline in edgeR.

### qRT-PCR

RNA was isolated from primary rat OPC/OLs and mouse tissue with the RNeasy Mini Kit (74104, QIAGEN) and stored at -80°C. cDNA was generated with SuperScript III First-Strand Synthesis (18080400, Thermo Fisher) and stored at -20°C. qPCR was performed with PowerUP SYBR Green (A25742, Thermo Fisher) on a QuantStudio 6 Flex Real-Time PCR System (4485691, Thermo Fisher). RT-qPCT primers were designed on the Integrated DNA Technologies (IDT) PrimerQuest program.

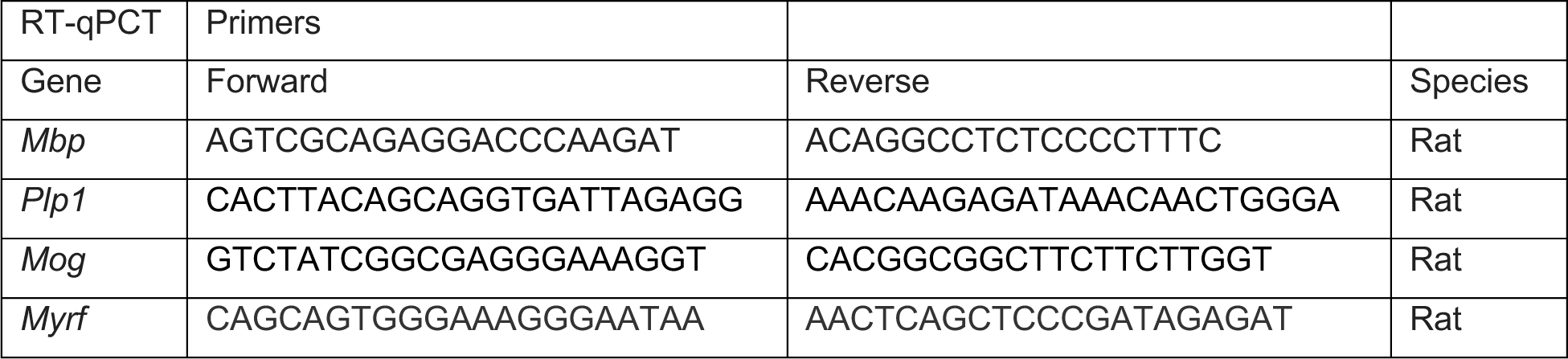
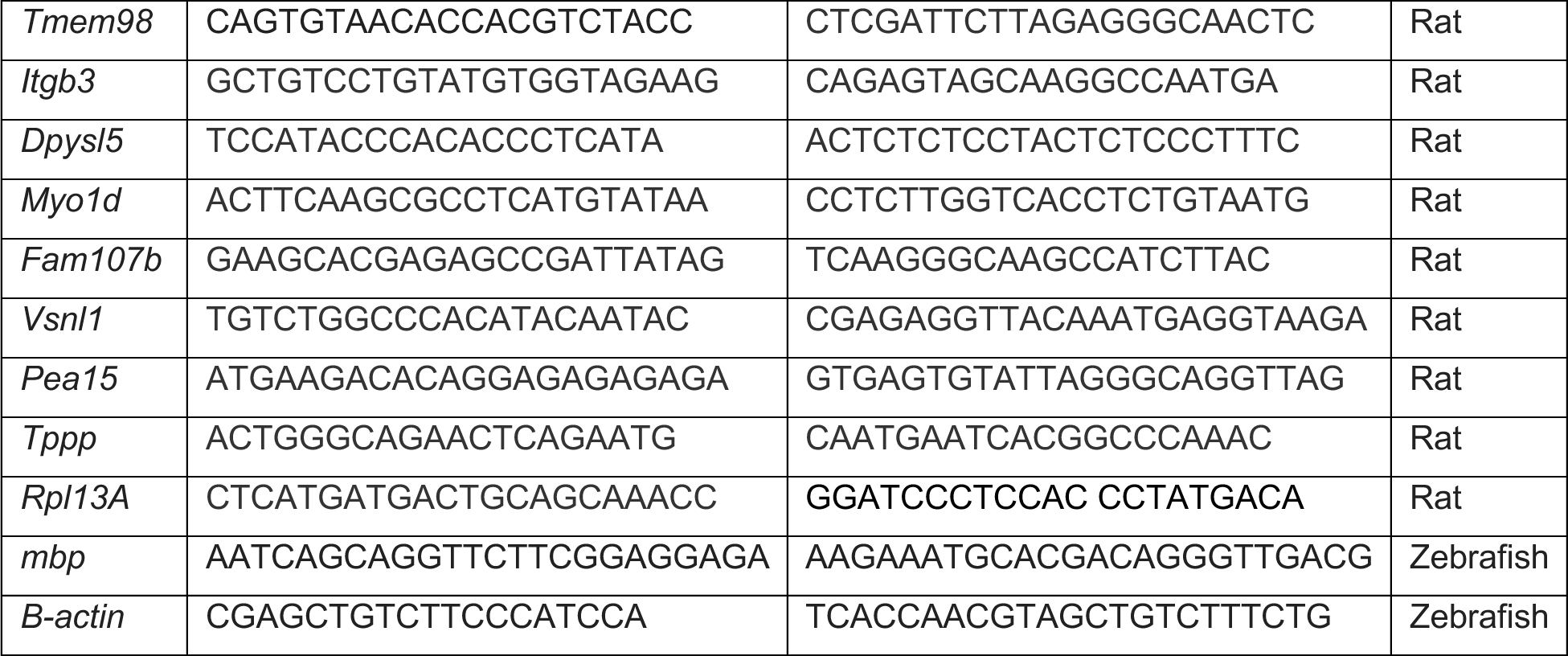

### Quantification and statistical analysis

Statistical analyses were conducted with Prism 10 (Graphpad). In all cases the figure legend indicates the statistical test used and p-values are presented in figures. Sample size is stated in figure legends. Animals were assigned to group based on genotype by random selection and analysis was conducted blinded to genotype.

## Data availability

Source data and accession numbers for proteomics datasets will be provided at the time of publication.

## Acknowledgements

We would like to thank current and past members of the Emery and Monks labs, particularly Suhail Akram, Emma Brennan, Austin Forbes, Tia Perry, and Adriana Reyes for excellent animal care. We would especially like to thank Dr. Breanne Harty of the Monk lab for her work on first characterizing FBXW7 in the zebrafish CNS and the mouse PNS. We would also like to thank Dr. Ronald Waclaw at Cincinnati Children’s Hospital for alerting us to the anti-MYRF antibody. We would like to thank Dr. Ashok Reddy of the OHSU Proteomic Core for his comments and suggestions while designing our proteomic experiments. This project was supported by the National Multiple Sclerosis Society (RG-1901-33272 to B.E. and K.R.M), the National Institutes of Health - National Institute of Neurological Disorders and Stroke (F31NS122433 to H.Y.C.). B.E. was supported by an endowment from the Warren family.

## Author contributions

H.Y.C, T.S., K.R.M, and B.E. conceived of the project. H.Y.C designed, performed, and analyzed all experiments with the following exceptions: J.L generated the *vo86* zebrafish line (H.Y.C performed all validation experiments and analyses). J.L performed and analyzed the *fbxw7* cell-specific CRISPR–Cas9-mediated gene disruption in Fig. 1f-h. R.A.D processed and imaged optic nerve TEM in Fig. 3. J.E.E and M.E.M. generated the *myrf^eu70^*zebrafish line in Fig. 6, and D.A.L. provided these mutants prior to publication. H.Y.C, K.R.M, and B.E. wrote the manuscript. All authors provided feedback on the manuscript and approved the submitted version.

## Materials and correspondence

Fish lines and reagents generated in this study including plasmids will be made available on request to the corresponding authors.

---

**Supplemental Fig. 1.**
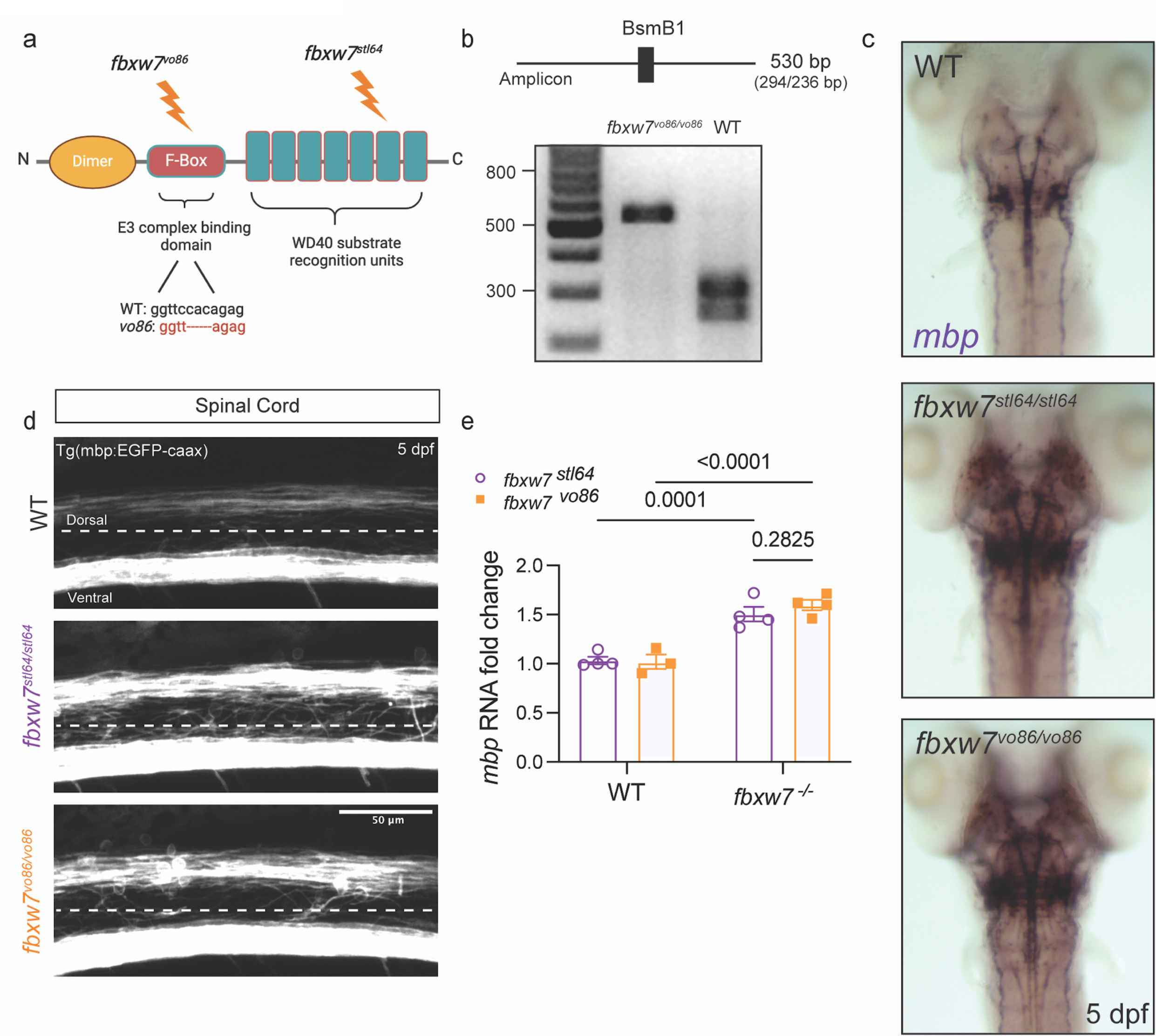
Creation and validation of *fbxw7^vo86^* zebrafish mutant allele. **a** Diagram of FBXW7 functional domains showing the locations of the previous ENU-induced mutation (*fbxw7^stl64^*) and new CRISPR-Cas9 generated mutation (*fbxw7^vo86^*). **b** Diagram of BsmB1 restriction enzyme digestion of a 560 bp amplicon of *fbxw7^vo86^* allele. Example of *fbxw7^vo86^* 560 bp genotyping PCR digested with BsmB1 restriction enzyme run on a 2% gel for WT and *fbxw7^vo86/vo86^* zebrafish larvae. **c** Representative images of *in situ* hybridization for *mbp* in 5 dpf WT, *fbxw7^stl64/stl64^,* and *fbxw7^vo86/vo86^* zebrafish larvae. **d** Representative images of the spinal cord from *fbxw7^stl64/stl64^* and *fbxw7^vo86/vo86^* Tg(mbp:EGFP-caax) zebrafish lines showing an increase in *mbp:EGFP-caax* intensity in both mutant alleles compared to WT control. **e** qRT-PCR for *mbp* from *fbxw7^stl64/stl64^*and *fbxw7^vo86/vo86^* whole zebrafish larvae relative to wild-type controls. Average ± SEM, WT controls N = 3, *fbxw7^-/-^* N=4, biological replicates (larvae). Statistical significance determined by two-way ANOVA. Created in BioRender. Emery, B. (2024) BioRender.com/l98g401.

**Supplemental Fig. 2.**
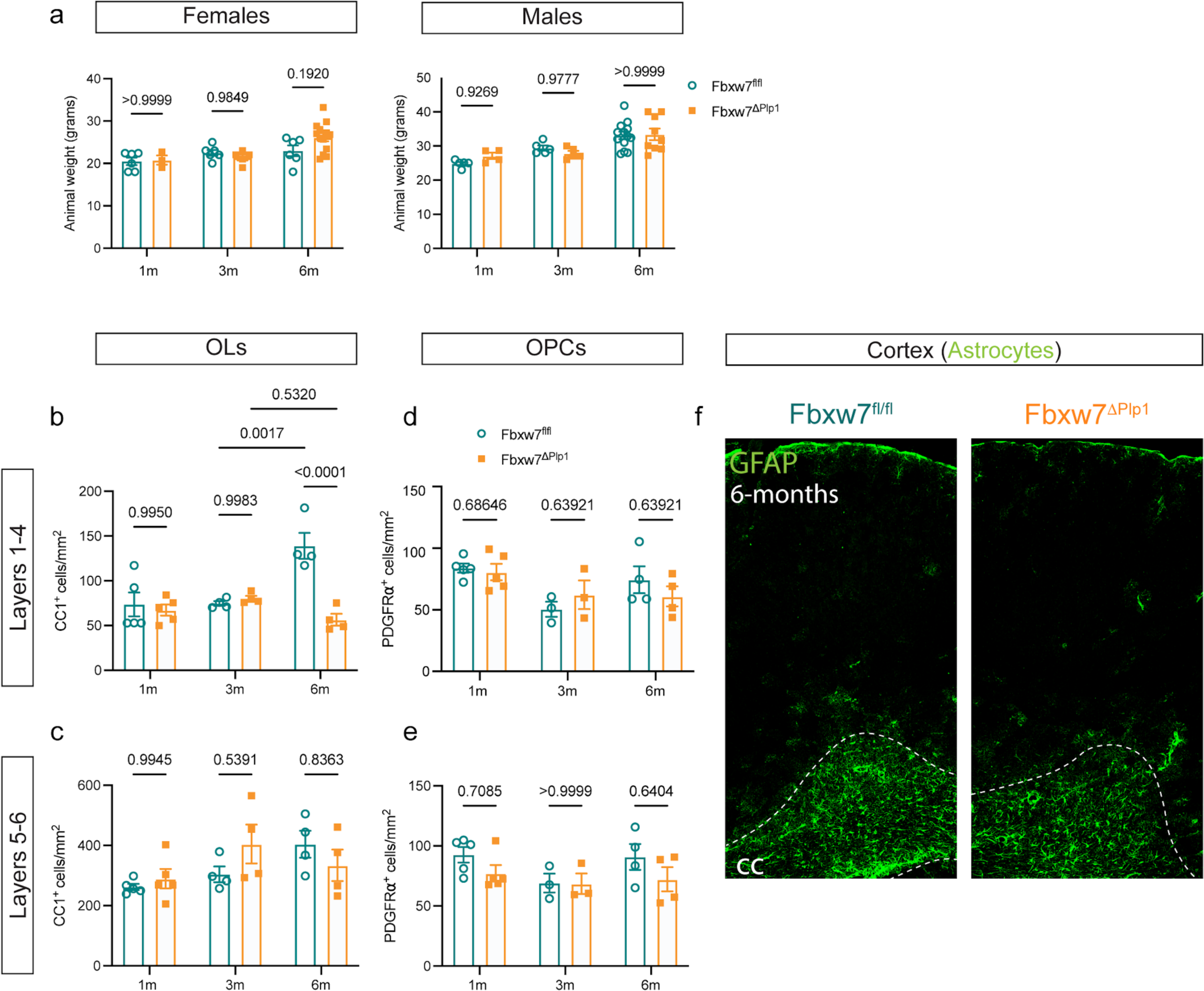
Fbxw7^fl/fl^; Plp1-CreERT mouse line characterization. **a** Animal weights for both male and female Fbxw7^fl/fl^ and Fbxw7^ΔPlp1^ mice at 1, 3, and 6 months post-TAM. **b** Quantification of OL number in cortical layers 1-4 and **c** 5-6. Average ± SEM, statistical significance determined by two-way ANOVA. **d** Quantification of OPC number in cortical layers 1-4, and **e** 5-6 at 1-month (Fbxw7^fl/fl^ N=4, Fbxw7^ΔPlp1^ N=4), 3 month (Fbxw7^fl/fl^ N=3, Fbxw7^ΔPlp1^ N=3), and 6 months post-TAM (Fbxw7^fl/fl^ N=4, Fbxw7^ΔPlp1^ N=4). **f** Representative images of cortical GFAP IF at 6 months post-TAM in Fbxw7^fl/fl^ and Fbxw7^ΔPlp1^. White dotted line delineates the border between the corpus callosum (CC) and cortex.

**Supplemental Fig. 3.**
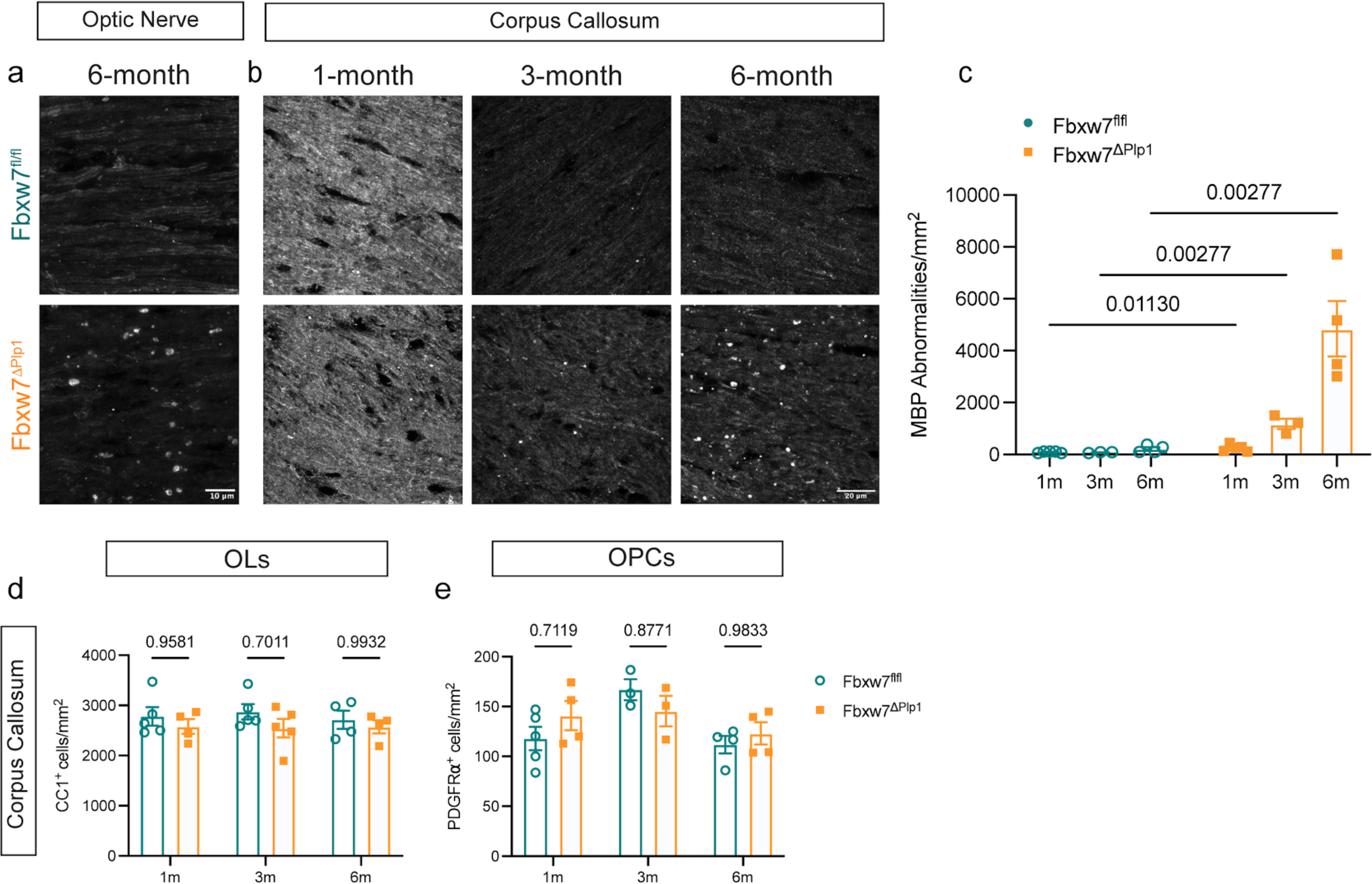
Deletion of *Fbxw7* results in myelin abnormalities in the corpus callosum and optic nerve. **a** Representative images of MBP IF at 6-months post-TAM in the optic nerve. **b** Representative images of MBP IF at 1, 3, and 6 months post-TAM in the corpus callosum. **c** Quantification of the density of MBP abnormalities in the corpus callosum at 1 month (Fbxw7^fl/fl^ N=4, Fbxw7^ΔPlp1^ N=4), 3 month (Fbxw7^fl/fl^ N=3, Fbxw7^ΔPlp1^ N=3), and 6 months post-TAM (Fbxw7^fl/fl^ N=4, Fbxw7^ΔPlp1^ N=4). Average ± SEM, statistical significance determined by two-way ANOVA. **d** OL densities in the corpus callosum in Fbxw7^fl/fl^ and Fbxw7^ΔPlp1^ mice at 1 month (Fbxw7^fl/fl^ N=5, Fbxw7^ΔPlp1^ N=4), 3 month (Fbxw7^fl/fl^ N=5, Fbxw7^ΔPlp1^ N=5), and 6 months post-TAM (Fbxw7^fl/fl^ N=4, Fbxw7^ΔPlp1^ N=4). Average ± SEM, statistical significance determined by two-way ANOVA. **e** Quantification of OPC densities in the corpus callosum at 1 month (Fbxw7^fl/fl^ N=5, Fbxw7^ΔPlp1^ N=4), 3-month (Fbxw7^fl/fl^ N=3, Fbxw7^ΔPlp1^ N=3), and 6 months post-TAM (Fbxw7^fl/fl^ N=4, Fbxw7^ΔPlp1^ N=4).

**Supplemental Fig. 4.**
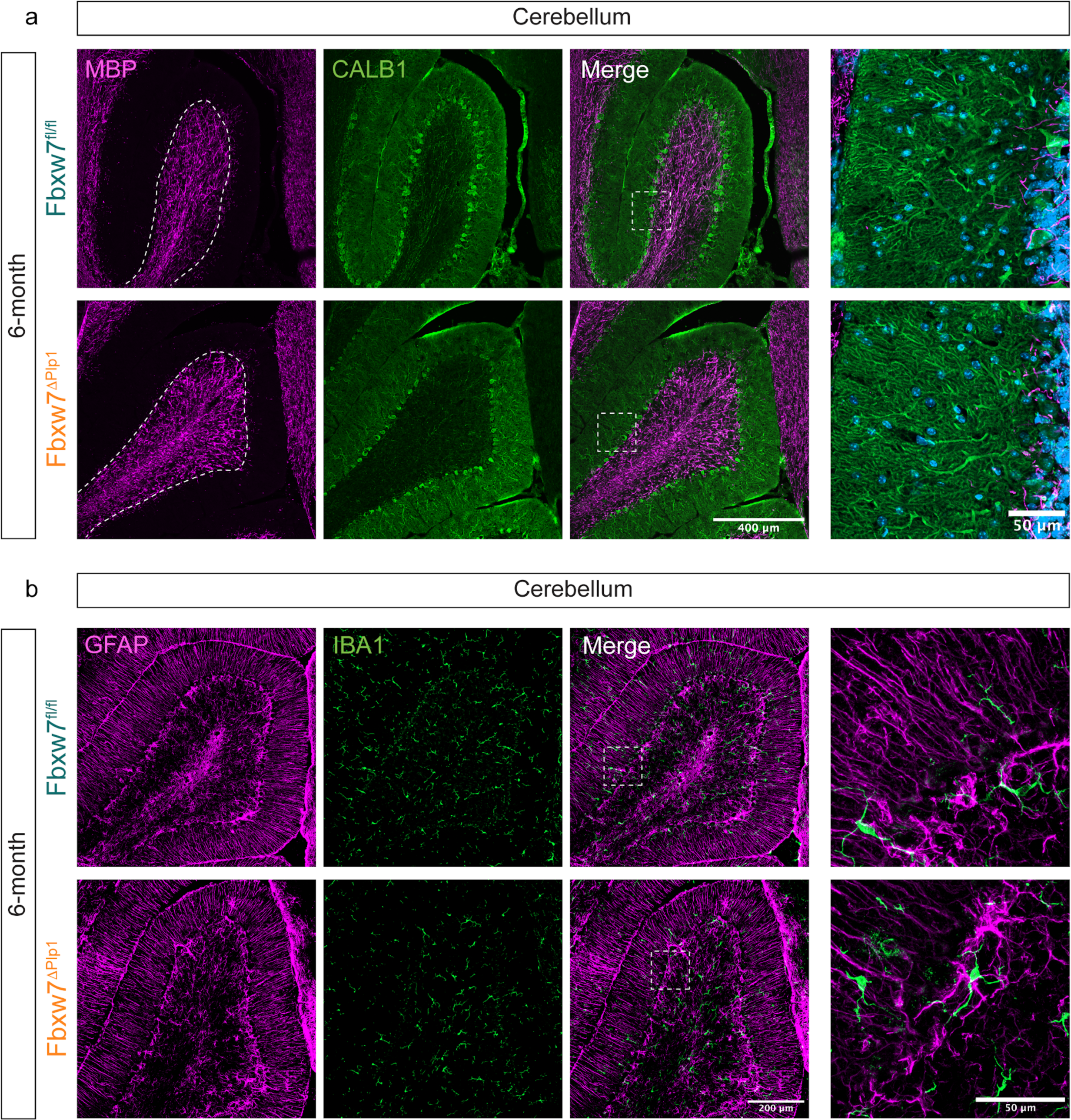
Fbxw7^ΔPlp1^ mice have normal cerebellar organization at 6-months post-TAM. **a** Representative images of MBP and Calbindin1 (CALB1) IF in the cerebellum of Fbxw7^fl/fl^ and Fbxw7^ΔPlp1^ mice at 6 months post-TAM. High resolution images of Calbindin1^+^ Purkinje cells show intact morphology in Fbxw7^ΔPlp1^ in the molecular layer of the cerebellum 6 months post-TAM. **b** Representative images of GFAP and IBA1 IF in the cerebellum of Fbxw7^fl/fl^ and Fbxw7^ΔPlp1^ mice at 6 months post-TAM.

**Supplemental Fig. 5.**
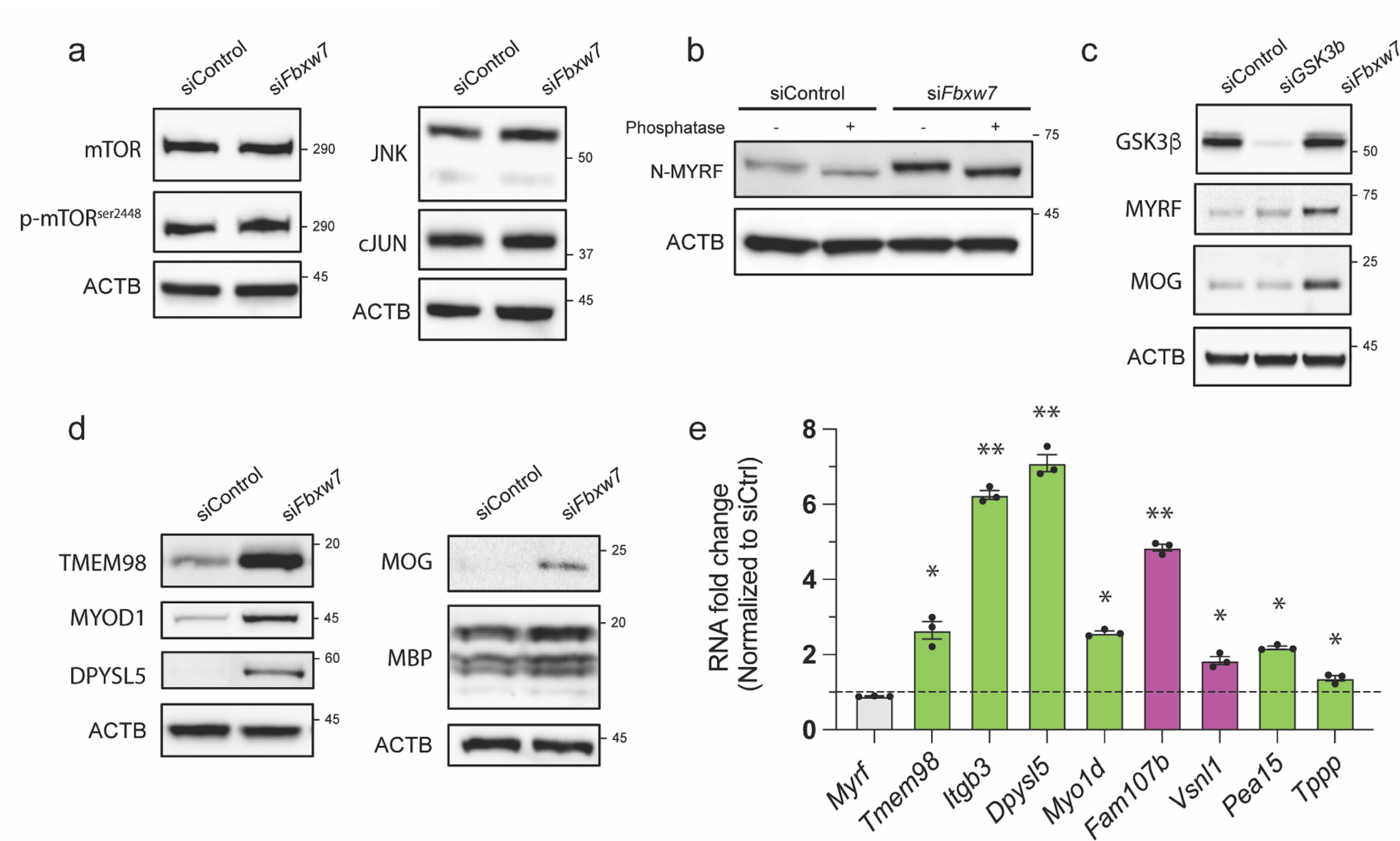
Western blot and RT-qPCR assessment of potential FBXW7 targets in OLs. **a** Western blots for candidate FBXW7 targets mTOR, p-mTOR^ser2448^, JNK and cJun in siControl and si*Fbxw7* electroporated rat OLs at 72 hours differentiation. **b** Western blot analysis for N-MYRF in siControl and si*Fbxw7* electroporated rat OL lysates with lambda phosphatase treatment of the lysates to reveal phosphorylation-dependent changes in molecular weight. **c** Western blot analysis of GSK3β, N-MYRF, and MOG in lysates from siControl, si*Gsk3b* and si*Fbxw7* electroporated rat OLs at 3 days differentiation. **d** Western blot analysis for top LC-MS hits TMEM98, MYOD1, DPYSL5 as well as myelin proteins MOG and MBP in siControl and si*Fbxw7* electroporated rat OLs at 72 hours differentiation. ACTB serves as a loading control. **e** qRT-PCR analysis of corresponding transcripts for enriched proteins from si*Fbxw7* electroporated OLs relative to siControl (* p < 0.05, ** p < 0.005). Average ± SEM, N = 3 (independent isolations) with 2 technical replicates (qPCR). Statistical significance determined by multiple unpaired, two-tailed Student’s t test.

**Supplemental Fig. 6.**
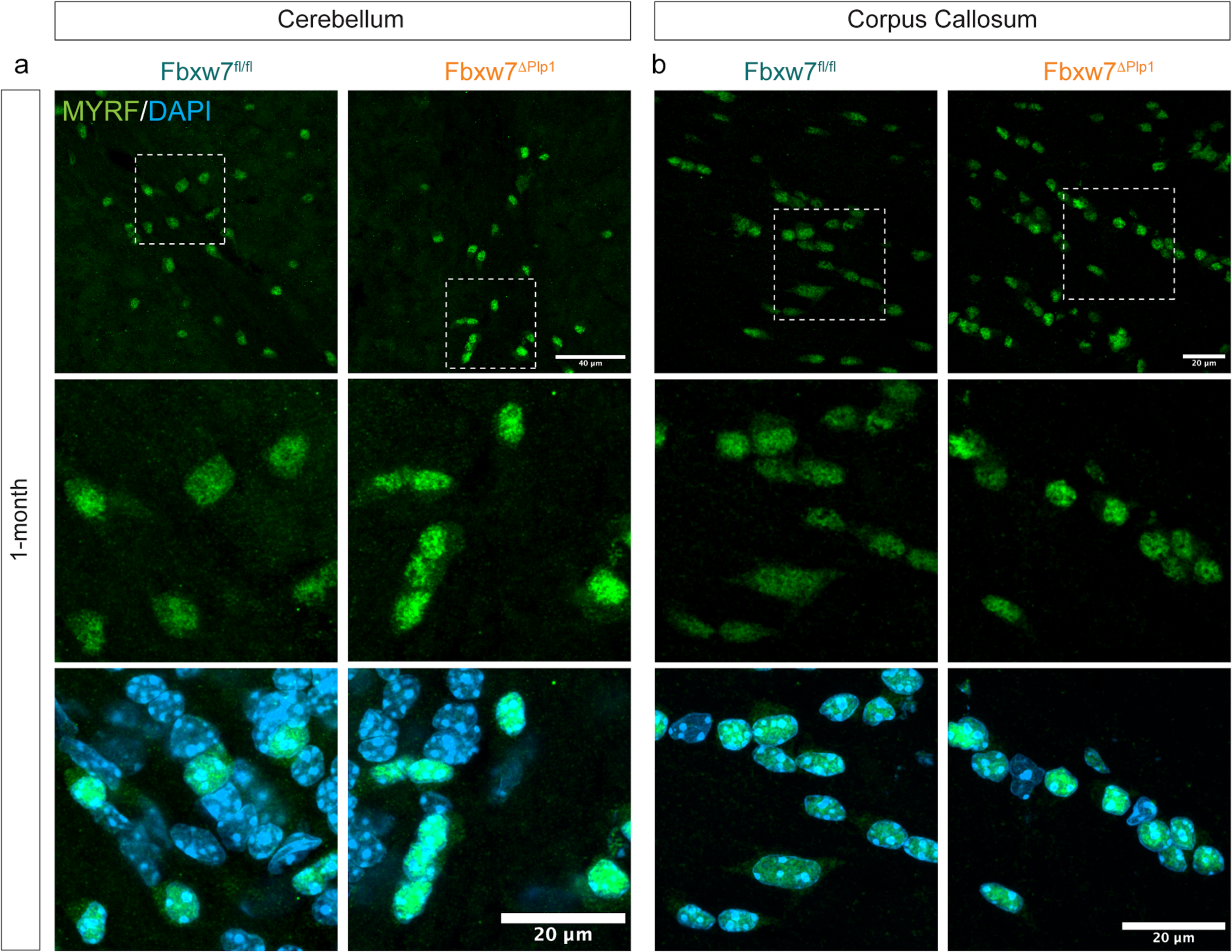
Deletion of *Fbxw7* in mature OLs results in increase nuclear MYRF in cerebellum and corpus callosum. **a** Representative images of MYRF IF in cerebellar white matter of Fbxw7^fl/fl^ and Fbxw7^ΔPlp1^ mice at 1 month post-TAM. **b** Representative images of MYRF IF in the corpus callosum of Fbxw7^fl/fl^ and Fbxw7^ΔPlp1^ mice at 1 month post-TAM.

## References

1. Waxman, S. G. & Bennett, M. V. L. Relative Conduction Velocities of Small Myelinated and Non-myelinated Fibres in the Central Nervous System. Nature. New Biol. 238, 217–219 (1972).

2. Ritchie, J. M. Physiological Basis of Conduction in Myelinated Nerve Fibers. in Myelin (ed. Morell, P.) 117–145 (Springer US, Boston, MA, 1984). doi:10.1007/978-1-4757-1830-0_4.

3. Nave, K.-A. & Trapp, B. D. Axon-Glial Signaling and the Glial Support of Axon Function. Annu. Rev. Neurosci. 31, 535–561 (2008).

4. Huxley, A. F. & Stämpfli, R. Direct determination of membrane resting potential and action potential in single myelinated nerve fibres. J. Physiol. 112, 476–495 (1951).

5. Czopka, T., ffrench-Constant, C. & Lyons, D. A. Individual Oligodendrocytes Have Only a Few Hours in which to Generate New Myelin Sheaths In Vivo. Dev. Cell 25, 599–609 (2013).

6. Zhou, Q., Choi, G. & Anderson, D. J. The bHLH Transcription Factor Olig2 Promotes Oligodendrocyte Differentiation in Collaboration with Nkx2.2. Neuron 31, 791–807 (2001).

7. Hughes, E. G., Kang, S. H., Fukaya, M. & Bergles, D. E. Oligodendrocyte progenitors balance growth with self-repulsion to achieve homeostasis in the adult brain. Nat. Neurosci. 16, 668– 676 (2013).

8. Hughes, E. G., Orthmann-Murphy, J. L., Langseth, A. J. & Bergles, D. E. Myelin remodeling through experience-dependent oligodendrogenesis in the adult somatosensory cortex. Nat. Neurosci. 21, 696–706 (2018).

9. Saher, G. et al. High cholesterol level is essential for myelin membrane growth. Nat. Neurosci. 8, 468–475 (2005).

10. Nawaz, S. et al. Actin filament turnover drives leading edge growth during myelin sheath formation in the central nervous system. Dev. Cell 34, 139–151 (2015).

11. Zuchero, J. B. et al. CNS Myelin Wrapping Is Driven by Actin Disassembly. Dev. Cell 34, 152– 167 (2015).

12. Cahoy, J. D. et al. A transcriptome database for astrocytes, neurons, and oligodendrocytes: a new resource for understanding brain development and function. J. Neurosci. Off. J. Soc. Neurosci. 28, 264–278 (2008).

13. Emery, B. et al. Myelin Gene Regulatory Factor Is a Critical Transcriptional Regulator Required for CNS Myelination. Cell 138, 172–185 (2009).

14. Koenning, M. et al. Myelin Gene Regulatory Factor Is Required for Maintenance of Myelin and Mature Oligodendrocyte Identity in the Adult CNS. J. Neurosci. 32, 12528–12542 (2012).

15. Elbaz, B. & Popko, B. Molecular Control of Oligodendrocyte Development. Trends Neurosci. 42, 263–277 (2019).

16. Marques, S. et al. Oligodendrocyte heterogeneity in the mouse juvenile and adult central nervous system. Science 352, 1326–1329 (2016).

17. Yeung, M. et al. Dynamics of oligodendrocyte generation and myelination in the human brain. Cell 159, (2014).

18. Sabri, M. I., Bone, A. H. & Davison, A. N. Turnover of myelin and other structural proteins in the developing rat brain. Biochem. J. 142, 499–507 (1974).

19. Saher, G. & Simons, M. Cholesterol and Myelin Biogenesis. in *Cholesterol Binding and Cholesterol Transport Proteins: Structure and Function in Health and Disease* (ed. Harris, J. R.) 489–508 (Springer Netherlands, Dordrecht, 2010). doi:10.1007/978-90-481-8622-8_18.

20. Tripathi, R. B. et al. Remarkable Stability of Myelinating Oligodendrocytes in Mice. Cell Rep. 21, 316–323 (2017).

21. Auer, F., Vagionitis, S. & Czopka, T. Evidence for Myelin Sheath Remodeling in the CNS Revealed by In Vivo Imaging. Curr. Biol. 28, 549-559.e3 (2018).

22. Hill, R. A., Li, A. M. & Grutzendler, J. Lifelong cortical myelin plasticity and age-related degeneration in the live mammalian brain. Nat. Neurosci. 21, 683–695 (2018).

23. Snyder, J. L., Kearns, C. A. & Appel, B. Fbxw7 regulates Notch to control specification of neural precursors for oligodendrocyte fate. Neural Develop. 7, 15 (2012).

24. Kearns, C. A., Ravanelli, A. M., Cooper, K. & Appel, B. Fbxw7 Limits Myelination by Inhibiting mTOR Signaling. J. Neurosci. 35, 14861–14871 (2015).

25. Harty, B. L. et al. Myelinating Schwann cells ensheath multiple axons in the absence of E3 ligase component Fbxw7. Nat. Commun. 10, 2976 (2019).

26. Sanchez, N. E. et al. Whole Genome Sequencing-Based Mapping and Candidate Identification of Mutations from Fixed Zebrafish Tissue. G3GenesGenomesGenetics 7, 3415–3425 (2017).

27. Nateri, A. S., Riera-Sans, L., Costa, C. D. & Behrens, A. The Ubiquitin Ligase SCFFbw7 Antagonizes Apoptotic JNK Signaling. Science 303, 1374–1378 (2004).

28. Ye, X. et al. Recognition of Phosphodegron Motifs in Human Cyclin E by the SCFFbw7 Ubiquitin Ligase*. J. Biol. Chem. 279, 50110–50119 (2004).

29. Yada, M. et al. Phosphorylation-dependent degradation of c-Myc is mediated by the F-box protein Fbw7. EMBO J. 23, 2116–2125 (2004).

30. Nakayama, S., Yumimoto, K., Kawamura, A. & Nakayama, K. I. Degradation of the endoplasmic reticulum-anchored transcription factor MyRF by the ubiquitin ligase SCFFbxw7 in a manner dependent on the kinase GSK-3. J. Biol. Chem. 293, 5705–5714 (2018).

31. Thompson, B. J. et al. Control of hematopoietic stem cell quiescence by the E3 ubiquitin ligase Fbw7. J. Exp. Med. 205, 1395–1408 (2008).

32. Li, J., Miramontes, T. G., Czopka, T. & Monk, K. R. Synaptic input and Ca2+ activity in zebrafish oligodendrocyte precursor cells contribute to myelin sheath formation. Nat. Neurosci. 27, 219–231 (2024).

33. Doerflinger, N. H., Macklin, W. B. & Popko, B. Inducible site-specific recombination in myelinating cells. Genes. N. Y. N 2000 35, 63–72 (2003).

34. Orthmann-Murphy, J. et al. Remyelination alters the pattern of myelin in the cerebral cortex. eLife 9, e56621 (2020).

35. Call, C. L. & Bergles, D. E. Cortical neurons exhibit diverse myelination patterns that scale between mouse brain regions and regenerate after demyelination. Nat. Commun. 12, 4767 (2021).

36. Almeida, R. G. The Rules of Attraction in Central Nervous System Myelination. Front. Cell. Neurosci. 12, (2018).

37. Klingseisen, A. et al. Oligodendrocyte Neurofascin Independently Regulates Both Myelin Targeting and Sheath Growth in the CNS. Dev. Cell 51, 730–744.e6 (2019).

38. Redmond, S. A. et al. Somatodendritic Expression of JAM2 Inhibits Oligodendrocyte Myelination. Neuron 91, 824–836 (2016).

39. Barron, T., Saifetiarova, J., Bhat, M. A. & Kim, J. H. Myelination of Purkinje axons is critical for resilient synaptic transmission in the deep cerebellar nucleus. Sci. Rep. 8, 1022 (2018).

40. Chung, S.-H., Guo, F., Jiang, P., Pleasure, D. E. & Deng, W. Olig2/Plp-positive progenitor cells give rise to Bergmann glia in the cerebellum. Cell Death Dis. 4, e546 (2013).

41. Winston, J. T., Koepp, D. M., Zhu, C., Elledge, S. J. & Harper, J. W. A family of mammalian F-box proteins. Curr. Biol. 9, 1180–S3 (1999).

42. Zhang, J. et al. Rack1 protects N-terminal phosphorylated c-Jun from Fbw7-mediated degradation. Oncogene 31, 1835–1844 (2012).

43. Richter, K. T., Kschonsak, Y. T., Vodicska, B. & Hoffmann, I. FBXO45-MYCBP2 regulates mitotic cell fate by targeting FBXW7 for degradation. Cell Death Differ. 27, 758–772 (2020).

44. Hornig, J. et al. The Transcription Factors Sox10 and Myrf Define an Essential Regulatory Network Module in Differentiating Oligodendrocytes. PLOS Genet. 9, e1003907 (2013).

45. Bujalka, H. et al. MYRF Is a Membrane-Associated Transcription Factor That Autoproteolytically Cleaves to Directly Activate Myelin Genes. PLOS Biol. 11, e1001625 (2013).

46. Li, Z., Park, Y. & Marcotte, E. M. A Bacteriophage Tailspike Domain Promotes Self-Cleavage of a Human Membrane-Bound Transcription Factor, the Myelin Regulatory Factor MYRF. PLOS Biol. 11, e1001624 (2013).

47. Aprato, J. et al. Myrf guides target gene selection of transcription factor Sox10 during oligodendroglial development. Nucleic Acids Res. 48, 1254–1270 (2020).

48. Jin, J., Ang, X. L., Shirogane, T. & Wade Harper, J. Identification of Substrates for F-Box Proteins. in Methods in Enzymology vol. 399 287–309 (Academic Press, 2005).

49. Madden, M. E. et al. CNS Hypomyelination Disrupts Axonal Conduction and Behavior in Larval Zebrafish. J. Neurosci. Off. J. Soc. Neurosci. 41, 9099–9111 (2021).

50. Gibson, E. M. et al. Neuronal Activity Promotes Oligodendrogenesis and Adaptive Myelination in the Mammalian Brain. Science 344, 1252304 (2014).

51. Hines, J. H., Ravanelli, A. M., Schwindt, R., Scott, E. K. & Appel, B. Neuronal activity biases axon selection for myelination in vivo. Nat. Neurosci. 18, 683–689 (2015).

52. Almeida, R. G. et al. Myelination of Neuronal Cell Bodies when Myelin Supply Exceeds Axonal Demand. Curr. Biol. 28, 1296–1305.e5 (2018).

53. Katanov, C. et al. N-Wasp Regulates Oligodendrocyte Myelination. J. Neurosci. 40, 6103– 6111 (2020).

54. Rosenbluth, J. Redundant myelin sheaths and other ultrastructural features of the toad cerebellum. J. Cell Biol. 28, 73–93 (1966).

55. Jaegle, M. et al. The POU proteins Brn-2 and Oct-6 share important functions in Schwann cell development. Genes Dev. 17, 1380–1391 (2003).

56. Kipreos, E. T. & Pagano, M. The F-box protein family. Genome Biol. 1, reviews3002.1 (2000).

57. Elbaz, B. et al. Phosphorylation State of ZFP24 Controls Oligodendrocyte Differentiation. Cell Rep. 23, 2254–2263 (2018).

58. Kornfeld, S. F., Cummings, S. E., Fathi, S., Bonin, S. R. & Kothary, R. MiRNA-145-5p prevents differentiation of oligodendrocyte progenitor cells by regulating expression of myelin gene regulatory factor. J. Cell. Physiol. 236, 997–1012 (2021).

59. Garnai, S. J. et al. Variants in myelin regulatory factor (MYRF) cause autosomal dominant and syndromic nanophthalmos in humans and retinal degeneration in mice. PLOS Genet. 15, e1008130 (2019).

60. Huang, H. et al. Interactive Repression of MYRF Self-Cleavage and Activity in Oligodendrocyte Differentiation by TMEM98 Protein. J. Neurosci. 38, 9829–9839 (2018).

61. Djannatian, M. et al. Two adhesive systems cooperatively regulate axon ensheathment and myelin growth in the CNS. Nat. Commun. 10, 4794 (2019).

62. Almeida, R. G., Czopka, T., ffrench-Constant, C. & Lyons, D. A. Individual axons regulate the myelinating potential of single oligodendrocytes in vivo. Development 138, 4443–4450 (2011).

63. Kim, H. et al. Notch-regulated perineurium development from zebrafish spinal cord. Neurosci. Lett. 448, 240–244 (2008).

64. Karttunen, M. J., Czopka, T., Goedhart, M., Early, J. J. & Lyons, D. A. Regeneration of myelin sheaths of normal length and thickness in the zebrafish CNS correlates with growth of axons in caliber. PLOS ONE 12, e0178058 (2017).

65. Peri, F. & Nüsslein-Volhard, C. Live Imaging of Neuronal Degradation by Microglia Reveals a Role for v0-ATPase a1 in Phagosomal Fusion In Vivo. Cell 133, 916–927 (2008).

66. Labun, K. et al. CHOPCHOP v3: expanding the CRISPR web toolbox beyond genome editing. Nucleic Acids Res. 47, W171–W174 (2019).

67. Almeida, R. G. & Lyons, D. A. Intersectional Gene Expression in Zebrafish Using the Split KalTA4 System. Zebrafish 12, 377–386 (2015).

68. Dugas, J. C. & Emery, B. Purification of Oligodendrocyte Precursor Cells from Rat Cortices by Immunopanning. Cold Spring Harb. Protoc. 2013, pdb.prot070862 (2013).

